# The Joint Impact of Deleterious Mutations, Dominance, and Gene Flow on Linked Neutral Variation in Structured Populations

**DOI:** 10.64898/2026.05.15.724312

**Authors:** Pierre Lesturgie, Alexandre Blanckaert, Vitor C. Sousa

## Abstract

Most species are geographically structured, which leaves characteristic signatures in neutral genomic regions used to reconstruct demographic history. Yet, these signatures can be distorted by linked selection due to deleterious mutations. This can reduce genetic diversity through Background *Selection*, or increase diversity through *Associative Overdominance*. While their effects are well characterized in panmictic populations, they remain largely unexplored in structured populations. We investigated an Isolation-with-Migration model using simulations across a range of migration, selection, dominance, and recombination parameters. We quantified the effects of deleterious mutations on standard summary statistics by simulating either genotypes at neutral and deleterious mutations (*genotype*-based approach) or genealogies and deleterious mutations (*Ancestral Recombination Graph*-based approach). The latter recovered similar patterns, directly relating genomic patterns to underlying coalescent processes. For lower recombination rates, we found a Background Selection-driven regime for weakly codominant mutations, characterized by lower diversity and increased genetic differentiation. For recessive mutations, we identified an Associative Overdominance-driven regime, characterized by increased diversity and lower genetic differentiation. This is reflected in the genealogies: Background Selection shrunk coalescence times and led to lineage sorting topologies, while Associative Overdominance stretched coalescence times and led to incomplete lineage-sorting topologies. Migration strongly affected genetic differentiation and topology frequencies but had weaker effects on diversity and coalescence times. Our results provide evidence of a transition from Associative Overdominance to Background Selection for recessive deleterious mutations in an isolation-with-migration model. Importantly, these processes leave distinct signatures on genealogies, highlighting that *Ancestral Recombination Graph*-based approaches have the potential for inferring linked-selection in structured populations.

**Article summary:** Jointly inferring demography and selection from genomic data is challenging due to the effect of deleterious mutations on linked neutral regions. While this effect has been characterized in panmictic populations, little is known about this effect in structured populations. Here, we use simulations to investigate how deleterious mutations shape genomic variation in a structured model, comparing *genotype*-based and *Ancestral Recombination Graphs*-based approaches. We showed that Background Selection and Associative Overdominance leave distinct signatures on summary statistics and genealogies. Our results highlighted the need to consider the selection-demography interplay for genomic inference, and the potential of *Ancestral-Recombinant-Graph* to disentangle this interplay.

## Introduction

The evolutionary history of species is shaped by demographic and selective processes, which leave specific signatures across genomes. Because these signatures are widely used to reconstruct the evolutionary history of species, accurately modeling these processes is crucial for understanding how species are structured in space, how they persist and how they adapt locally across their range (Hoban et al. 2016; Tigano and Friesen 2016). Yet, because demographic history and selection interact and often generate confounding genomic patterns (Hoban et al. 2016), quantifying their respective contributions to genetic diversity remains challenging. Linked selection can bias demographic reconstructions (Ewing and Jensen 2016; Johri et al. 2021; Smith and Hahn 2024), but the reverse is also true, demographic history can bias the detection of selection (Jensen et al. 2005). Disentangling selective and demographic signatures at the genomic level requires a detailed understanding of how demographic and selective forces interact.

Most species are structured in meta-populations, i.e., set of demes connected by migration, rather than in a single panmictic (random mating) population, a common assumption when investigating selection and demography (e.g., Charlesworth et al. (1993); Charlesworth (2012); Zhao and Charlesworth (2016)). Yet such spatial structure is expected to generate genomic patterns distinct from those under panmixia (Hudson 1991; Wakeley 2009). Indeed, genome-wide patterns of diversity are commonly used to infer key demographic parameters, such as effective population size (*Ne*), effective migration rate (*M*), and times of population divergence (e.g., Bertorelle et al. (2010); Excoffier et al. (2021)). These approaches are based upon the neutral theory of evolution (Kimura 1968), i.e., shaped by genetic drift and demography. However, due to the impact of selection on linked neutral variation, many studies have questioned this common assumption (Charlesworth and Charlesworth 2018; Kern and Hahn 2018), since there is evidence for pervasive effects of linked selection in many animal and plant taxa (Sella et al. 2009; Cutter and Payseur 2013; Pouyet et al. 2018). Linked selection arises when the increase in frequency of beneficial alleles (Barton 2000) or the removal of deleterious ones (Charlesworth et al. 1993; Charlesworth 2012) alters patterns of genetic diversity at nearby linked neutral sites. The extent of this is determined by the strength of selection and recombination (Charlesworth et al. 1993; Charlesworth et al. 1997; Charlesworth 2012).

Deleterious mutations, by affecting cellular processes and physiology, have a negative impact on fitness irrespective of the environment (Agrawal and Whitlock 2012). Hence, deleterious mutations can have major consequences on the evolution of populations, for example leading to inbreeding depression especially in small and/or structured populations (Charlesworth and Charlesworth 1999; Whitlock 2002; Charlesworth and Willis 2009). The strength of the deleterious effect is given by the selective coefficient (*s*) and the degree of dominance (*h*), which relate to fitness (*w*) at a given biallelic locus as follows: *w*=1 for homozygotes without deleterious alleles, *w*=1-*hs* for heterozygotes, and *w*=1-*s* for homozygotes for the deleterious allele. In panmictic populations, two selective regimes, resulting from selection affecting deleterious mutations, have been described: one driven by Background Selection (BGS) and a second one by Associative Overdominance (AOD). BGS refers to the reduction of neutral genetic diversity in regions linked to deleterious mutations compared to the amount expected under neutrality (Charlesworth et al. 1993; Charlesworth 2012; Zhao and Charlesworth 2016; Cvijović et al. 2018). Typically, a BGS-driven regime arises for codominant or slightly recessive mutations with moderate fitness effects (Nordborg et al. 1996), as documented in Pouyet et al. (2018) and Gilbert et al. (2020). In the case of slightly deleterious and recessive mutations, arguably the most common type of deleterious mutation (Peters et al. 2003; Shaw and Chang 2006; Agrawal and Whitlock 2011), an alternative regime driven by AOD may emerge. AOD is a form of balancing selection that occurs when different haplotypes accumulate complementary sets of recessive deleterious mutations, leading to higher fitness of heterozygotes compared to homozygotes, as recessive mutations are (partially) hidden from selection in heterozygous individuals (Ohta and Kimura 1970; Ohta 1971a; Ohta 1971b; Zhao and Charlesworth 2016; Becher et al. 2020; Gilbert et al. 2020). Empirical examples supporting AOD are scarce (see Butlin and Day (1985); Jay et al. (2021); Waller (2021)), but consistent patterns have been found in regions of low recombination in *Drosophila* (Mukai and Yamaguchi 1974; Albornoz and Dominguez 1994; Schou et al. 2017; Becher et al. 2020) and humans (e.g., Gilbert et al. (2020)). Due to the advantage of heterozygotes driven by the coexistence and maintenance of complementary haplotypes, AOD is expected to result in higher levels of genetic diversity than expected under neutrality. This contrasts with expected lower levels of genetic diversity resulting from BGS. Despite this dichotomy, the two processes are connected by two important features. (1) Both AOD and BGS are expected to leave stronger signatures in regions of low recombination, where the lack of recombination limits the persistence of deleterious mutations-free haplotypes and amplifies their effect on linked neutral mutations. (2) When mutations are recessive, a transition between AOD and BGS occurs: as the deleterious effect (*s*) increases, a regime driven by AOD shifts into a regime driven by BGS (Zhao and Charlesworth 2016; Gilbert et al. 2020). AOD and BGS can thus be considered as bounds of a continuum, with effects on linked neutral variation negatively correlated with the recombination rate.

The effects of deleterious mutations in structured populations remain poorly characterized (but see Charlesworth et al. (1997); Zeng and Corcoran (2015); Ewing and Jensen (2016); Aeschbacher et al. (2017)). Theoretical studies have focused on codominant deleterious mutations, predicting that when migration is sufficiently low or absent, BGS decreases local genetic diversity (*π*), increase genetic differentiation (*F_ST_*) (Charlesworth et al. 1997; Zeng and Corcoran 2015; Aeschbacher et al. 2017) and shifts the Site Frequency Spectrum (*SFS*) towards low-frequency variants, compared to neutral expectations (Zeng and Corcoran 2015). When migration increases, local *π* may still decrease (Hasan and Whitlock 2024), but *F_ST_* is mostly unaffected (Matthey-Doret and Whitlock 2019; Hasan and Whitlock 2024). Population structure may also influence the emergence of pseudo-overdominance, as crosses between populations carrying different fixed recessive deleterious mutations can generate pseudo-overdominance through their masking in heterozygotes (Waller 2021). The effect of recessive deleterious mutations that could lead to an AOD-driven regime, and its transition to the BGS-driven regime, have however not been investigated in structured populations.

This gap can now be addressed more directly through recent developments in inference of *Ancestral Recombination Graphs* (ARGs) from population genomics datasets, which are revolutionizing the study of natural populations (Lewanski et al. 2024; Nielsen et al. 2025), opening the door to study the interaction between demographic history, recombination and selection. ARGs describe the ancestry relations between sampled DNA sequences, accounting for coalescent and recombination events (Hudson 1991). Coalescent-based frameworks assuming neutrality are now widely used to infer complex demographic histories, (e.g., Kelleher and Lohse (2020); Excoffier et al. (2021)). Selection is expected to also affect topologies and branch lengths, and theoretical studies suggest that linked selection can be modeled using coalescent with altered parameters, e.g., BGS being represented as a reduction of *Ne* (Hudson and Kaplan 1988). To date, the genealogical consequences of linked selection have been primarily characterized for BGS in panmictic populations (Hudson and Kaplan 1995; Nordborg 1997; Barton et al. 2004; Zeng 2013; Torres et al. 2020; Johri et al. 2021) (but see Zeng and Corcoran (2015)). In consequence, predictions for the properties of gene trees jointly accounting for demography and linked selection caused by deleterious mutations remain largely unexplored, despite being required to interpret observed local genealogies from estimated ARGs.

Here we sought to (1) investigate the signatures of deleterious mutations on linked genetic variability in structured populations; and (2) determine how deleterious mutations affect linked neutral gene genealogies. To that end, we performed simulations under an isolation with migration model in which recombination, dominance, fitness effects, and migration can all vary. We showed that both AOD and BGS-driven regimes can emerge when recombination is low, and that there is a transition from AOD to BGS in the case of recessive mutations. While selection determines the impact of these two processes, the presence of gene flow modulates their signatures. The ARG-based approach allows better discrimination between demographic and selective processes by leveraging explicit genealogical properties. This highlights its potential for future inference of linked selection in structured populations.

## Methods

### Demographic scenario and genomic architecture

To investigate the effect of linked deleterious mutation on the pattern of diversity of structured populations, we considered an Isolation with Migration (IM) model (**Figure 1a**) in which, going forward in time, an ancestral population of size *N_ANC_*=1,000 diploids evolved for 10,000 generations (a burn-in to guarantee that the population reached mutation-selection-drift equilibrium (*t_eq_*>4*N_ANC_*)). At the end of this period, the population split into two populations of *N*=1,000 diploids individuals, with the newly formed populations remaining either isolated (*M=0*) or exchanging *M=*2*Nm*=2.5 migrants per generation for *T_DIV_*=1,000 generations. *M*=2.5 was chosen to represent a clear gene-flow regime, well above the classic 4*Nm*=1 at which migration is expected to substantially counteract differentiation by genetic drift. We modeled the genome (**Figure 1b**) as a single chromosome of 100,000 base pairs, where the first and last 10,000bp were regions where mutations were deleterious (with a negative fitness effect of 2*Ns*, and a dominance coefficient *h*), while mutations in the remaining 80,000 bp were neutral.

**Figure 1.**
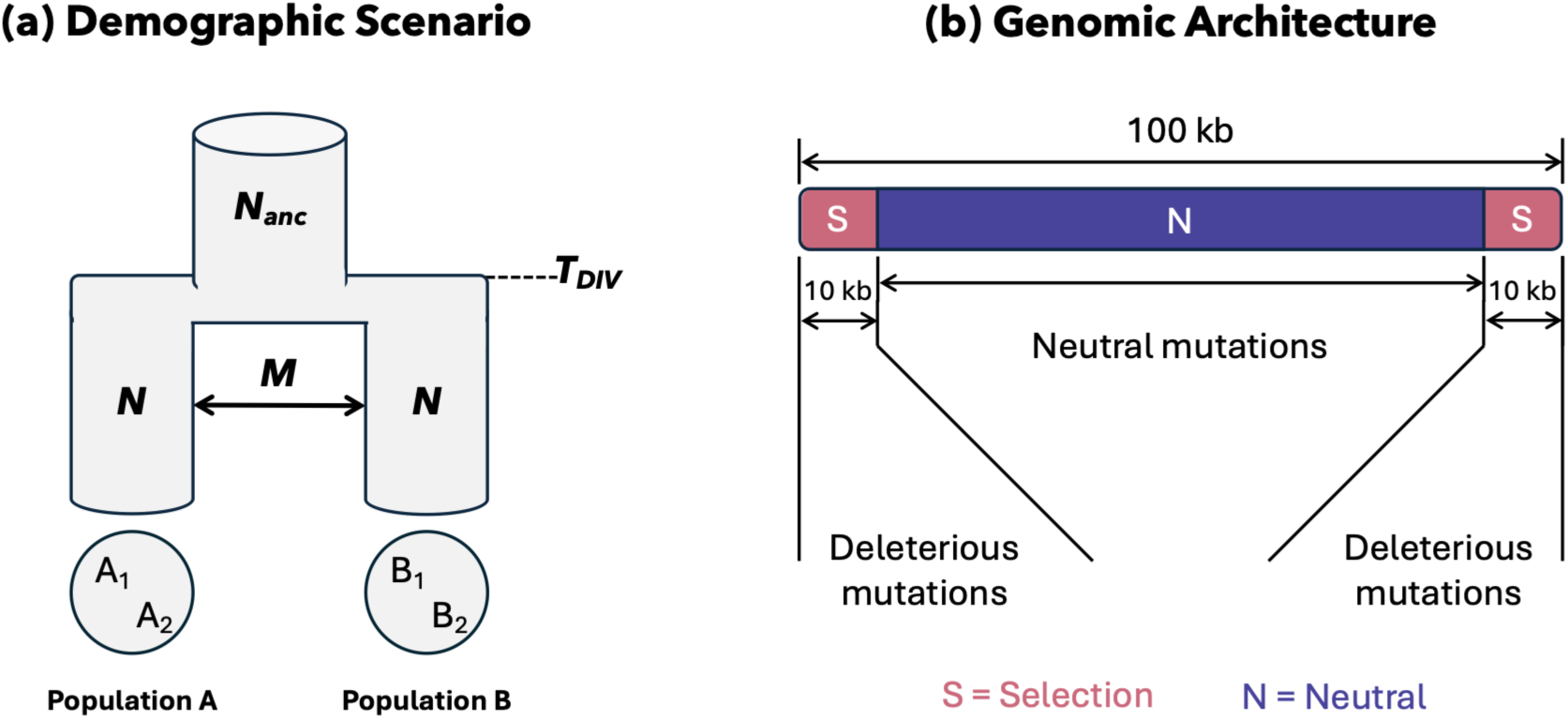
Demographic scenario (a) and genomic architecture (b). **(a)** Going forward in time, the demographic scenario depicts an ancestral population of size N=1,000 diploid individuals evolving for 10,000 generations until it splits into two sub-populations of size N and exchanging symmetric M migrants (either M=0 or M=2.5) per generation for 1,000 generations. A1 and A2 (respectively B1 and B2) represent two haploid lineages sampled from a diploid individual from population A (respectively population B) when using the ARG-based framework (see methods/results). **(b)** The genomic architecture consists of a single chromosome of length 100kb, divided into three regions. Mutations in the first and last 10kb are deleterious, while mutations in the central region (80kb) are neutral.

### Simulations

We performed a large set of simulations (**Table 1**) using *SLiM v5* (Haller et al. 2025), varying recombination rate (*r*), selective coefficients (2*Ns*), dominance (*h*) and migration (*M*=2*Nm*) with identical coefficient for the ancestral and the two descending populations (scripts for the pipeline available at https://github.com/PierreLesturgie/Pipeline-Slim-BGS-AOD). Three recombination rates were tested: *r*=2.5e-6, 2.5e-7 and 2.5e-8 per site per generation (Table 1), which corresponds to 0.1, 1 and 10 times the mutation rate, which was fixed to *µ*=2.5e-7 per site per generation. The mutation rate was chosen as a balance between computational efficiency and validity of the infinite site model, as more important than the absolute values of the parameters are their relationship. Thus, for the same value of *θ* = 4*Nμ*, we performed a pilot test of several combinations of mutation rates and effective sizes to ensure that patterns converged to the expectations under the infinite sites model. A mutation rate of 2.5e-7 per site per generation and an effective size of 1,000 individuals ensure both validity under the infinite site model and computational efficiency. We assumed that deleterious and neutral mutations all occurred at the same rate and investigated the case of identical selective and dominance coefficients for all deleterious mutations. Discrete values of deleterious selective coefficients are indicated in **Table 1**. Two dominance cases were investigated: deleterious mutations were either codominant (*h*=0.5) or recessive (*h* = 0.01). Finally, we investigated both a case of isolation (*M*=0) and of a symmetric migration between the two populations (*M* = 2.5). For each set of parameters, we performed 1024 simulations.

**Table 1.** Simulated parameter values.

| Parameter | Values | Note |
| --- | --- | --- |
| Recomb. Rate (R) | 2.5e-6, 2.5e-7, 2.5e-8 |  |
| Selection coefficient ($2N_s$) | 0, 0.02, 0.1, 0.2, 1, 2, 10, 20, 100, 200, 1000, 2000 | Also tested on a subset: $2N_s \sim \text{Exp}(1/20)$ and $2N_s$ drawn from a joint $2N_s h$ distribution |
| Dominance ($h$) | 0.01, 0.5 | |
| Migration ($M=2Nm$) | 0, 2.5 | Also tested on a subset: $M=0.5$ |

To further confirm the robustness of the results, we investigated two additional scenarios. (1) We considered a distribution of the deleterious effects in which 2*Ns* was drawn from an exponential distribution of mean 2*Ns*=20. (2) To mimic the correlation between *s* and *h* that have been estimated in empirical studies in model species (Phadnis and Fry 2005; Huber et al. 2018; Becher et al. 2020), we performed simulations where *s* and *h* were drawn from a joint distribution. To facilitate comparisons, we used a joint distribution such that the mean *s* was ∼0.01. We used a discretize exponential distribution, with bins centered around the discrete values of 2*Ns* mentioned in **Table 1**, with the dominance value deduced from the equation (1) from Huber et al. (2018), parametrized in such a way that *h*=0.5 for *s*=0 and *h*=0.01 for *s*=0.01.

In addition to these simulations, we also compared, for a subset of selection coefficients (2*Ns* of 20 and 200) the effect of deleterious mutations on *π_g_* in a single panmictic population, by computing the values for the ancestral population, and an intermediate migration rate *M*=0.5, for a subset of deleterious coefficients (2*Ns* = 0, 10, 20, 100, 200).

### Summary statistics

We computed a set of different summary statistics on the neutral (80,000 bp) region of the chromosome following two different approaches, either based on the actual genotype of the individuals (*genotype*-based approach) or based on the gene tree sequences (*ARG*-based approach).

For the *genotype*-based approach, at the end of the simulation, we sampled 10 diploid individuals per population. We computed a set of different statistics based on the extracted genotype of these 10 individuals: mean pairwise difference (*π*), absolute divergence (*d_xy_*), genetic differentiation (Hudson’s *F_ST_*) and the average Derived Allele Frequency per individual (*DAFi*).

The *ARG*-based approach is based on gene tree sequences subsampled for a sample of *n*=1 diploid per deme, as assumed in model-based inference methods (Laetsch et al. 2023). The expected value of all the summary statistics considered can be derived either considering a sample size of 2 lineages sampled from the same (*π_g_*), or different populations (*d_xy-g_, F_ST_*), or a sample size of 4 lineages (2 from each population, *DAF_i_*). Hence, for the ARG-based we used a sample of 1 diploid per population, such that we could derived expected values for all statistics, while also investigating the tree topologies and coalescent time distributions. While such a different sampling scheme is not ideal to directly compare the distributions of statistics of the ARG-based and the genotype-based approaches, the expected values should be similar as the sample size should impact the variance. To get the tree sequences, we tracked the entire genealogical trees of the individuals simulated. We computed summary statistics based on the output tree sequences using a coalescent framework based on *tskit* (Ralph et al. 2020; Wong et al. 2024) and *pyslim* (Gopalan et al. 2025). The first step consisted in sampling two diploid individuals, one from each deme. In the case of multiple roots, i.e., the genealogy did not trace back to a common ancestor, we used the *recapitation* method to infer the most common recent ancestor of the sample (Gopalan et al. 2025). For each tree sequence, we only kept the neutral trees from the full tree sequence and computed the following summary statistics from branch lengths and coalescence times: (1) *π_g_* (2\**E[T_within_],* where *T_within_* is the coalescent times for a pair of lineages sampled in the same population); *d_xy-g_* (2\**E[T_between_]* where *T_between_* is the coalescent times for a pair of lineages sampled from each population) and the Site Frequency Spectrum *(SFS)* were directly obtained from branch lengths and Hudson’s *F_ST_* indirectly through *π_g_* and *d_xy-g_*; (2) *DAF_i_* using the ratio between the average Time to the Most Recent Common Ancestor (*T_MRCA_)* and total branch length over the tree sequence (Pouyet et al. 2018); (3) distribution of topology and coalescence times from branch lengths and coalescence events (see below).

All ARGs can be classified into eight topologies named Topology 1 *INTRA (T1_INTRA_)* Topology 1 *INTER (T1_INTER_),* Topology 2 *INTRA (T2_INTRA_),* Topology 2 *INTER (T2_INTER_),* Topology 3 *INTRA* (*T3_INTRA_*), Topology 3 *INTER* (*T3_INTER_*), Topology 4 and Topology 5 (**Table 2**). Considering four sampled haploid genomes (A_1_ and A_2_ from population A; B_1_ and B_2_ from population B), Topology 1 is observed when the first and second coalescent events occur between external branches and Topology 2 happens when the first coalescent event occurs between two external branches and the second between the resulting internal branch and an external branch. For topology 1 and 2, we further distinguished between *INTRA* and *INTER* classification (**Table 2**). In details, Topology 1 *INTRA* (hereafter *T1_INTRA_*) and Topology 2 *INTRA* (*T2_INTRA_*) are characterized by the first coalescent event occurring between individuals from the same population. Oppositely, Topology 1 *INTER* (*T1_INTER_*), and Topology 2 *INTER* (*T2_INTER_*) are defined by the first coalescent event occurring between individuals from different populations. Topologies 3, 4 and 5 correspond to multiple merger coalescent topologies, involving three, three and four coalescence events at the same time, respectively. Note that Topology 3 is also distinguished into *INTRA* and *INTER* classification based on the first coalescence event occurring either between the same deme or between the two demes. Because Topologies 3 and 4 represented almost no trees and Topology 5 no trees, we only represented the results for Topologies 1 and 2. For each simulation we quantified (1) the distribution of topologies observed over the trees in the sequence, and the coalescence times (*T_4_* being the first coalescent event, *T_3_* the second and *T_2_* the last, i.e., the *T_MRCA_* in our case of four sampled lineages), conditioned on each topology. Finally, to have robust estimates of summary statistics, each estimate was repeated 10 times and averaged across the 10 runs. Scripts to estimate statistics based on gene tree sequences are available on GitHub (https://github.com/PierreLesturgie/GTS).

**Table 2.**
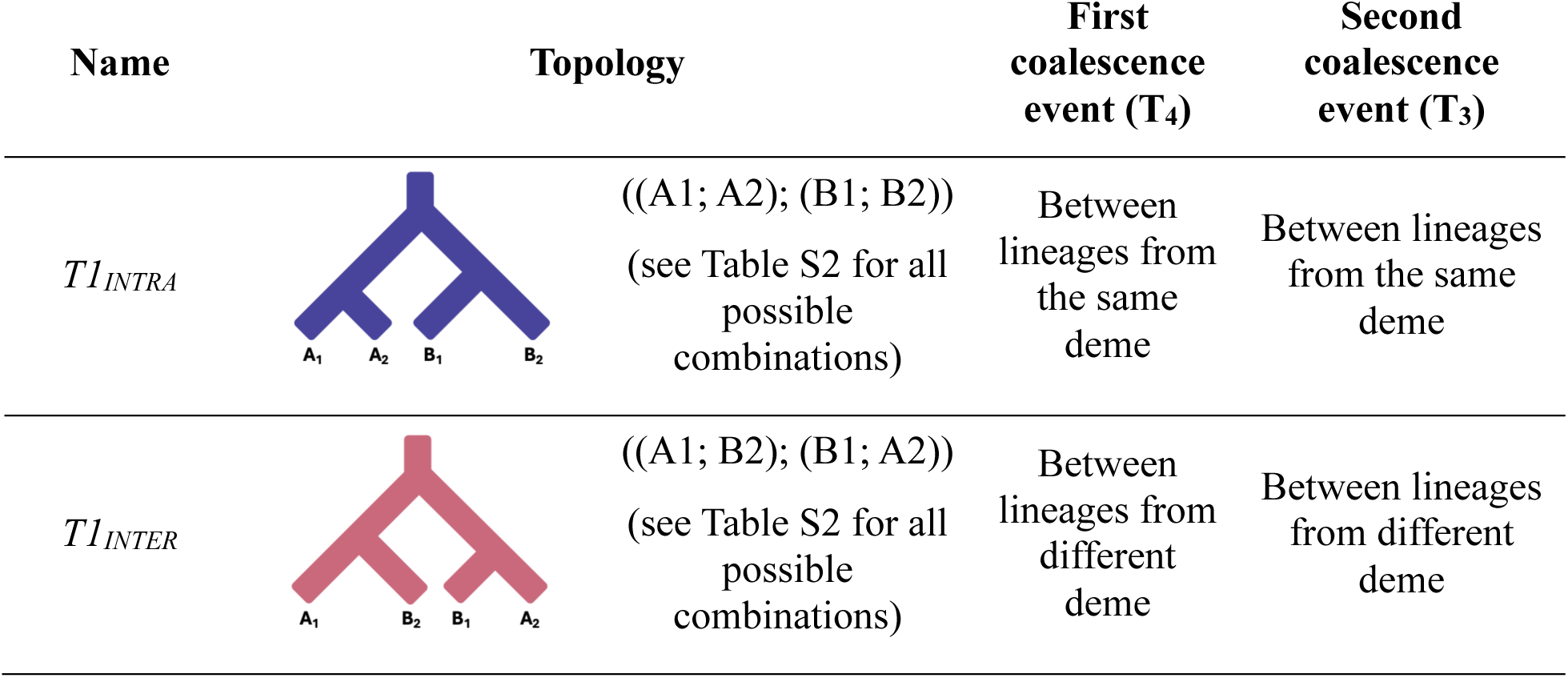
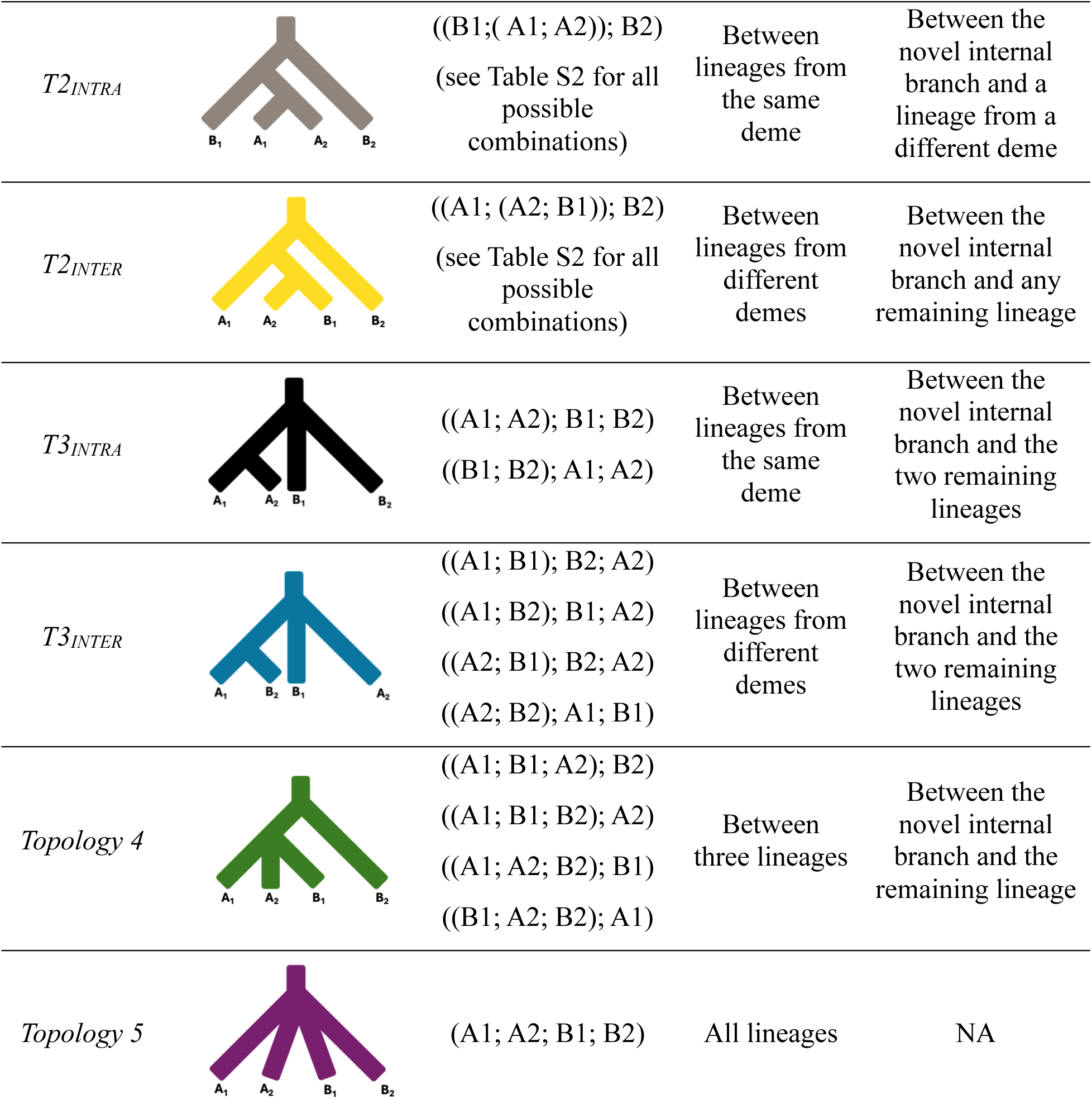
Classification of the six possible topologies with a sample of four haploids, sampling A1 and A2 from population A and B1 and B2 from population B. Topologies are first classified based on whether the first coalescence event occurs within demes (INTRA topologies). Topologies 1 correspond to cases where the first two coalescence events occur within or between demes; topologies 2 to cases where the second coalescence event involves the novel internal branch and a remaining lineage. Topologies 3, 4 and 5 represent multiple merger coalescent events, respectively involving three, three or four simultaneous lineages. Here T_4_ represents the first coalescence event, T_3_ the second.

Hereafter, we described deviations from neutrality as differences in summary statistics relative to the neutral expectation and, for each summary statistic (*θ*) we reported the ratio *b* between the extremum (maximum or minimum) median value obtained for simulations under selection (*θ_S_*) and the median value under neutrality (*θ_N_*), i.e., 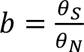 (**Table S1**).

## Results

In the following sections, we described in detail the results obtained with a recombination rate 10 times lower than the mutation rate (*r*=2.5e-8 per site and per generation), since it was the condition where the effects of linked selection were stronger (although patterns were slightly recovered for *r*=2.5e-7, **Figures S1-S5**).

### Codominant mutations in isolation decrease diversity and increase differentiation

We examined the effect of codominant deleterious mutations (*h* = 0.5) in the absence of migration (*M*=0), using the *genotype-based* approach (**Figure 2**). As expected, for slightly (*2Ns*<1) and strongly (2*Ns*>200) deleterious mutations, none of the four statistics (the mean pairwise difference (*π*), absolute divergence (*d_xy_*), genetic differentiation (*F_ST_*), and the averaged Derived Allele Frequency per individual (*DAF_i_*) deviated from neutrality. This was consistent with deleterious mutations behaving either neutrally or lethally. For intermediate value of 2*Ns* (∈ [1,200]) we recovered patterns consistent with BGS (Charlesworth et al. 1997; Zeng and Corcoran 2015; Aeschbacher et al. 2017): all four statistics reach their extreme value around 2*Ns*=20, with *π, d_xy_* and *DAF_i_* being minimized while *F_ST_* is maximized. From this point onward, we referred to the selective coefficients at which we found those patterns as the BGS-driven regime.

**Figure 2.**
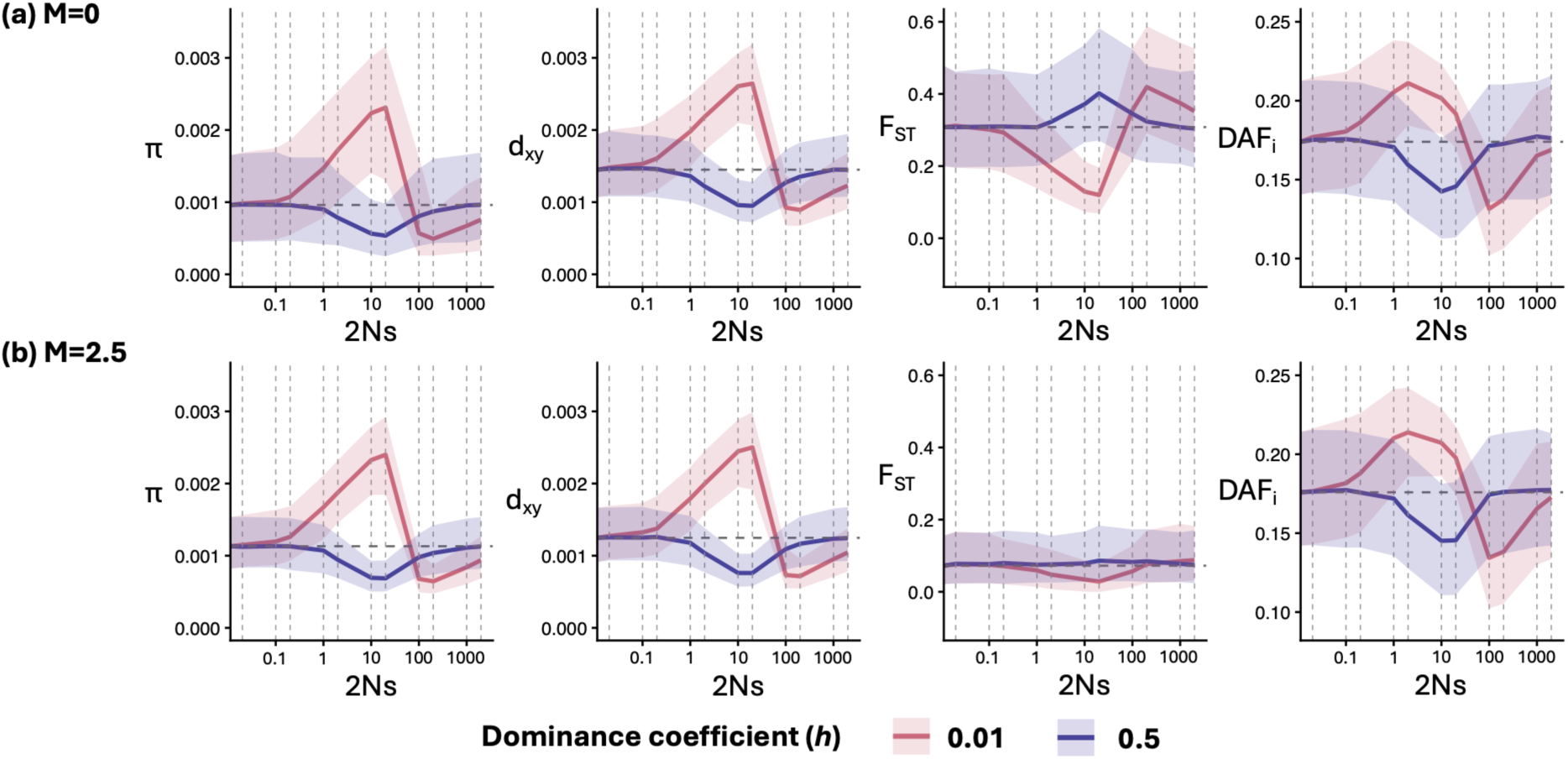
BGS and AOD generate different and unique patterns. We displayed here the effect of the deleterious fitness effect on summary statistics computed using the genotype-based approach under a recombination rate of r=2.5e-8 per site per generation in the absence **(a)** or presence **(b)** of migration. The summary statistics displayed are the mean pairwise difference (π), the absolute divergence (d_xy_), the genetic differentiation (F_ST_), and the average Derived Allele Frequency per individual (DAFi). For each statistic, the trajectory of the median value as a function of the selective coefficient (2Ns) is represented and the areas around the curve represent the 75% confidence interval computed over 1024 simulations. For each plot, the two curves represent codominant (h=0.5) mutations (blue) or recessive (h=0.01) mutations (red). The vertical grey lines represent the 2Ns values used in the simulations and the dashed horizontal line represent the median neutral value. The x-axis is displayed in log scale.

### Recessive mutations in isolation increase diversity and decrease differentiation

For recessive mutations (*h*=0.01), two regimes emerged (**Figure 2, Table S1**). First, as the deleterious effects of mutations increased, all statistics started to depart from neutrality for 2*Ns* ≥ 0.2. Three statistics (*π, d_xy_* and *DAF_i_*) increased to reach maximum values at 2*Ns*=2 (*DAFi*) or 2*Ns*=20 (*π*, *d_xy_*), (*b* = 2.40, 1.82 and 1.21, respectively); while *F_ST_* decreased to reach its minimum value at 2*Ns* = 20 (*b* = 0.39, **Table S1**). This increase of *π* and *DAF_i_* was consistent with AOD (Gilbert et al. 2020). From this point onwards, we interpreted these variations as the AOD-driven regime. After the peak in the AOD-driven regime, as the strength of selection increased, *π*, *d_xy_* and *DAF_i_* dropped (*b* = 0.51, 0.62 and 0.76, respectively) at 2*Ns* values of 100 (*DAFi*) and 200 (*π, d_xy_*), whereas *F_ST_* increased (*b*=1.36) at 2*Ns* = 200. This constituted a subsequent BGS-driven regime, consistent with an AOD-BGS transition (Gilbert et al. 2020). In contrast with codominant mutations, the statistics obtained in the presence of recessive mutations did not match the one obtained for neutrality, even at 2*Ns*=2000.

### Migration blurs signatures of genetic differentiation

Migration strongly affected the process of removal of mutations and thus the impact of BGS and AOD on genomic patterns (**Table S1, Figure 2**). Under *M*=2.5, *DAF_i_* remained similar to *M*=0, consistent with its expected robustness to demography (Pouyet et al. 2018). For codominant mutations, *DAF_i_* was reduced between 2*Ns* ∈ [1,200] (*b*=0.83 at 2*Ns*=10), consistent with the BGS-driven regime. For recessive mutations, *DAFi* increased between 2*Ns* ∈ [0.2,20] (*b*=1.22 at 2*Ns*=2), consistent with the AOD-driven regime, before decreasing for 2*Ns*≥100 (*b*=0.76 at 2*Ns*=100), reflecting the AOD-BGS transition. Both *π* and *d_xy_* behaved qualitatively similarly in the presence or absence of migration: we observed a decrease for 2*Ns* ∈ [1,200] for codominant mutations, and for 2*Ns* ≥ 100 for recessive mutations (BGS-driven regime) and an increase for 2*Ns* ∈ [0.2,20] for recessive mutations (AOD-driven regime) (**Figure 2)**. With migration, *F_ST_* increased under BGS (*b* = 1.19 at 2*Ns*=20 for *h*=0.5, and *b*=1.21 at 2*Ns*=2000 for *h*=0.01), but remained close to the expected value under neutrality overall, lacking the pronounced maximums observed in isolation (**Figure 2**). Yet the pronounced decrease of *F_ST_* in the AOD-driven regime was preserved for 2*Ns* ∈ [1,100] (*b* = 0.39 at 2*Ns*=20, **Figure 2**, **Table S1**).

### The ARG-based approach consistently recovers the effects of BGS and AOD

As expected, the *ARG*-*based* approach recovered patterns consistent with the *genotype-based* approach (**Figure 3, Table S1**). Under isolation and for codominant mutations, all four statistics (*π_g_*, *d_xy-g_*, *DAF_i_* and *F_ST_*) showed patterns consistent with a BGS-driven regime over similar 2*Ns* ranges (2*Ns* ∈ [1,100]) and reached similar *b* values to those obtained with the *genotype-based* approach. The Site Frequency Spectrum (*SFS*) also reflected the BGS-driven regime for 2*Ns* ∈ [1,100], with a strong reduction in the proportion of shared polymorphism between populations (*b* = 0.46, 2*Ns*=20), and a slight increase in the proportion of private polymorphism (i.e., singletons restricted to one population, *b* = 1.07, 2*Ns*=10). For recessive mutations, *π_g_, d_xy-g_, DAFi* and *F_ST_* all showed patterns consistent with the AOD-driven regime for 2*Ns* ∈ [0.2,20], followed by a transition to the BGS-driven regime at 2*Ns* = 100. The *SFS* also reflected this transition: for 2*Ns* ∈ [0.2,20], the proportion of shared polymorphism strongly increased (*b*=2.28 at 2*Ns*=10) and the proportion of private polymorphism slightly decreased for 2*Ns* ∈ [0.2,10] (*b*=0.93 at 2*Ns*=1), consistent with the AOD-driven regime. For 2*Ns*≥100, the proportion of shared polymorphism strongly decreased (*b*=0.41, 2*Ns*=200) and the proportion of private polymorphism increased (*b*=1.12, 2*Ns*=100), consistent with the BGS-driven regime. With migration, patterns remained comparable to those obtained with the *genotype-based* approach (**Figure 3, Table S1**). All summary statistics *π_g_, d_xy-g_, F_ST_* and *DAFi* showed the same trend, and the *SFS* retained signatures of AOD and BGS across similar 2*Ns* ranges, though with reduced magnitude in comparison to isolation.

**Figure 3.**
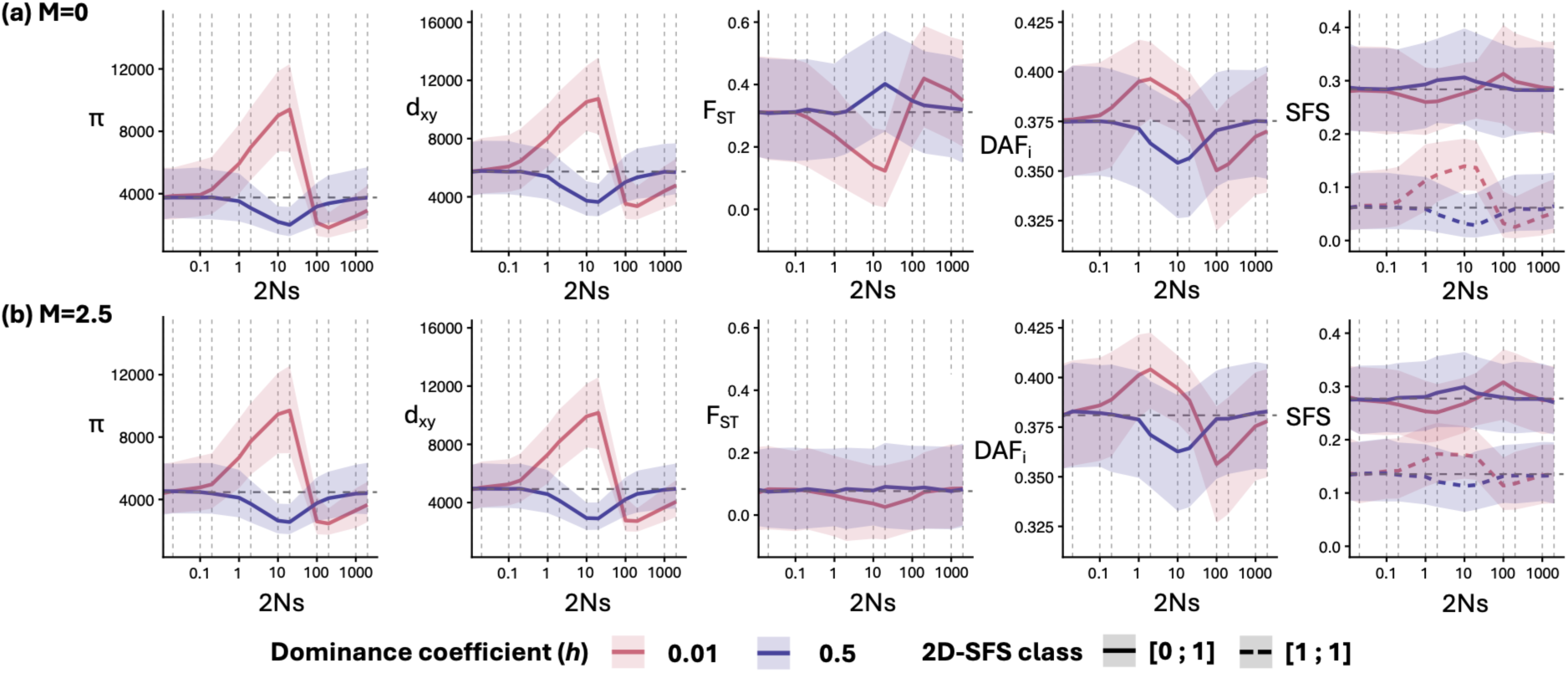
The unique patterns of AOD and BGS are similarly captured with the ARG approach. We displayed here the effect of the deleterious fitness effect on summary statistics computed using the ARG-based approach under a recombination rate of r=2.5e-8 per site per generation in the absence **(a)** or presence **(b)** of migration. The summary statistics displayed are the mean pairwise difference in branch lengths (π_g_), the absolute divergence in branch lengths (d_xy-g_), the genetic differentiation (F_ST_), the average Derived Allele Frequency per individual (DAFi) and the private (full line) or shared (dashed line) singletons classes of the two-dimensional Site Frequency Spectrum (2D-SFS). The π_g_ and d_xy-g_ are computed in branch lengths in generations and respectively represent 2*E[T_within_] and 2*E[T_between_], where T_within_ and T_between_ are the coalescent times for a pair of lineages sampled in the same population, or one from each population, respectively. For each statistic, the trajectory of the median value as a function of the selective coefficient (2Ns) is represented and the areas around the curve represent the 75% confidence interval computed over 1024 simulations. For each plot, the two curves represent codominant (h=0.5) mutations (blue) or recessive (h=0.01) mutations (red). The vertical grey lines represent the 2Ns values used in the simulations and the dashed horizontal line represents the median neutral value. The x-axis is displayed in log scale.

### The joint effect of deleterious mutations and gene flow determines topology distribution

Under neutrality and in isolation, *INTRA* topologies were the most common (mean=75%: 40% *T2intra*, 35% *T1intra*), indicating that in at least one of the populations, the two lineages coalesced more recently with each other than with a lineage from the other population; whereas *INTER* topologies (mean=24%) were mostly generated by Topology 2 (16% *T2inter*, 8% *T1inter*), the remaining 1% being multiple merger topologies. Thus, under neutrality and in isolation, only 35% of gene trees were consistent with the population tree and lineage sorting (*T1intra* topology). These results matched theoretical expectations (see Supplementary material). When including selection, the topology distributions differed greatly from neutrality and in opposite ways for the BGS and AOD-driven regimes. For codominant mutations (**Figure 4b, Table S1**), the BGS-driven regime (2*Ns* ∈ [2, 2000]) was characterized by an increase in the proportion of *T1_INTRA_* (*b*=1.42, 2*Ns* = 20), a slight decrease in *T2_INTRA_* (*b*=0.94, 2*Ns*=20), and a strong reduction in INTER topologies (*b* = 0.48 for *T1_INTER_* and 0.50 for *T2_INTER_* at 2*Ns* = 20). This change in the distribution of topologies indicated that, in the BGS-driven regime, the chance of lineage sorting occurring increased, i.e., gene trees and population trees were more likely to match. For recessive mutations (*h*=0.01), the patterns at *2Ns* values consistent with BGS-driven regime closely matched that of *h*=0.5 mutations for *2Ns*≥100. For 2*Ns* values consistent with the AOD-driven regime, we found increased incomplete lineage sorting for 2*Ns* ∈ [0.2, 20]): the proportion of *INTRA* topologies decreased (*T1_INTRA_*: *b*=0.51; *T2_INTRA_*: *b*=0.86 at *2Ns*=20), and *INTER* topologies strongly increased (*T1_INTER_*: *b* = 2.15, *T2_INTER_*: *b*= 1.88 at *2Ns*=20).

**Figure 4.**
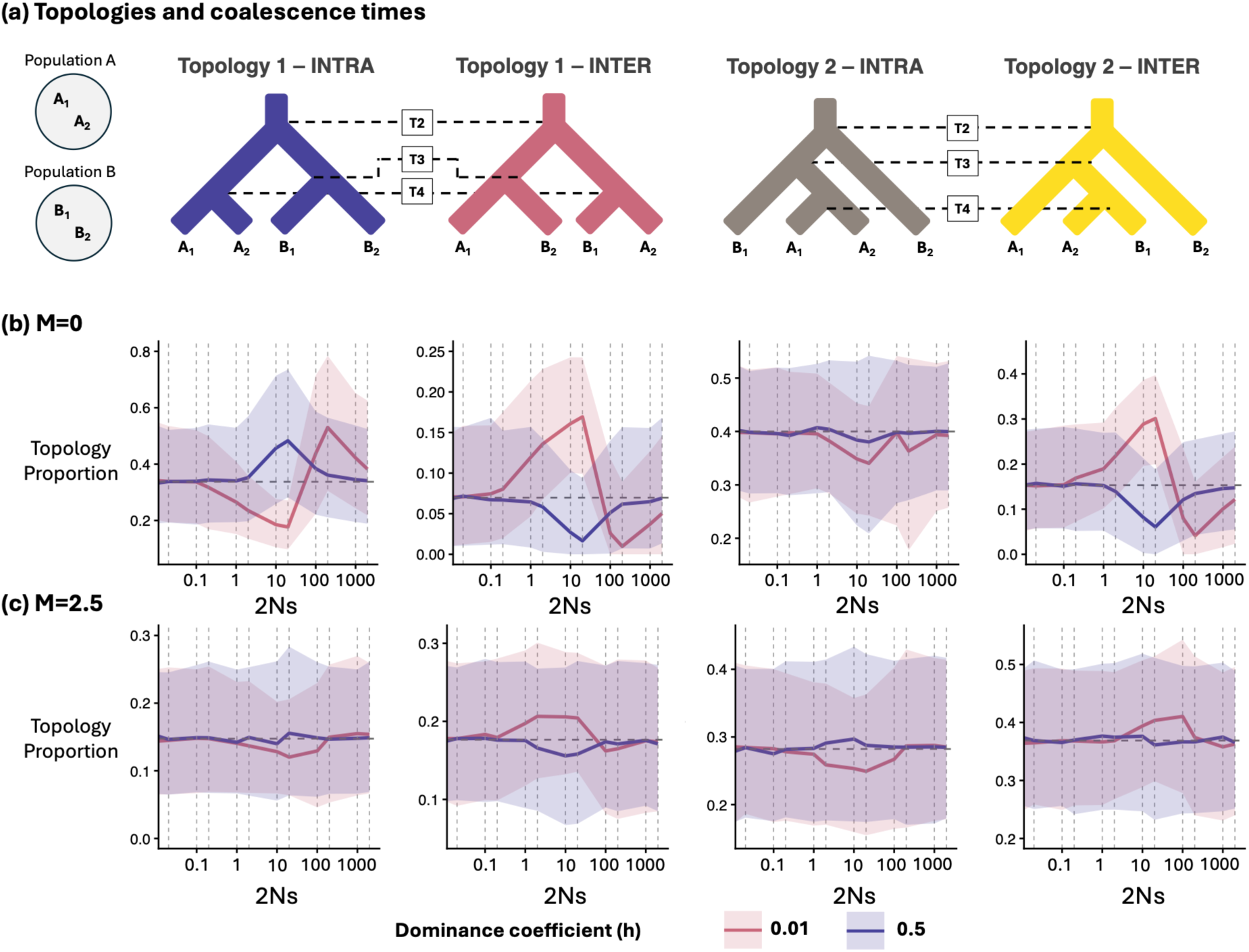
AOD and BGS distort the distribution of tree topologies. **(a)** We displayed here the four possible topologies in the tree sequence (excluding multiple mergers). Considering four haploid genomes (A1 and A2 in population A; B1 and B2 in population B), Topology 1 is observed when the first and second coalescent events occur between external branches and Topology 2 happens when the first coalescent event occurs between two external branches and the second between the resulting internal branch and an external branch. Two subsets are devised: Topology 1 INTRA (T1_INTRA_) and Topology 2 INTRA (T2_INTRA_) occur when the first coalescent occurs between genomes from the same population. Topology 1 INTER (T1_INTER_), and Topology 2 INTER (T2_INTER_) occur when the first coalescent occurs between genomes from different populations. **(b)** and **(c)**: proportion of the topology respectively to the topology displayed above each plot, in the context of isolation **(b)** and migration **(c)**. For each proportion of topology, the trajectory of the median value as a function of the selective coefficient (2Ns) is represented and the areas around the curve represent the 75% confidence interval computed over 1024 simulations. For each plot, the two curves represent codominant (h=0.5) mutations (blue) or recessive (h=0.01) mutations (red). The vertical grey lines represent the 2Ns values used in the simulations and the dashed horizontal line represents the median neutral value. The x-axis is displayed in log scale.

With migration (*M*=2.5) and under neutrality, we observed, as expected, more *INTER* topologies (55%) than in isolation. The rarest topology was *T1_INTRA_* (16%), indicating that only a small proportion of gene trees had reciprocal monophyly, i.e., complete lineage sorting. In the BGS-driven regime, the distribution of tree topologies was weakly affected by migration (**Table S1, Figure 4c**). The only consistent signal was a mild decrease of *T1_INTER_* for 2*Ns* ∈ [1, 100] for codominance (*b*=0.90 at 2*Ns*=10), and for 2*Ns* ∈ [100, 200] for recessive mutations (*b*=0.92 at 2*Ns*=100). However, AOD signals remained detectable, albeit weaker than in isolation: the proportion of *T1_INTRA_* and *T2_INTRA_* decreased for 2*Ns* ∈ [2, 200] (*b*=0.84 and 0.88 at 2*Ns*=20), while *T1_INTER_* increased for values of 2*Ns* ∈ [1, 20] (*b*=1.14), and *T2_INTER_* increased by for 2*Ns* ∈ [10, 100] (*b*=1.12).

### Coalescence times are stretched under AOD and shrunk under BGS irrespective of gene flow

The purging of deleterious mutations altered coalescence times in opposite directions for the BGS and AOD-driven regimes, with different impact of migration between the two regimes (**Figure 5**). In general, compared to neutrality, BGS lead to reduced coalescent times (*b*<1), whereas AOD leads to increased coalescent times (*b*>1, Table S1). Moreover, under the BGS-driven regime, two key patterns arose: (1) scaling factors (*b*) were nearly identical for *h* = 0.5 and *h* = 0.01 mutations, although the values of 2*Ns* at which the extreme effect occur differed between the two dominance scenarios, and (2) in the presence or absence of migration, compared to neutrality, all coalescent times were similarly affected in all topologies. Under the BGS-driven regime, coalescent times were always reduced by at least 15% indicating that *b*<0.85. In the presence of migration (*M*=2.5), all scaling factors were similar for the different topologies (**Table S1**). Under the AOD-driven regime, all coalescence times peaked at 2*Ns*=20 in the presence or absence of migration, and: (1) migration only slightly altered the shifts in coalescent times for certain topologies; and (2) on average, *T_2_* and *T_3_* increased more than *T_4_*, consistently with stronger increases in *d_xy_* than in *π*, although this was less pronounced for *M*=2.5 (**Table S1**). Overall, compared to neutrality, AOD lead to an increase in coalescent times of at least 28% (i.e., *b*>1.28) (**Table S1**).

**Figure 5.**
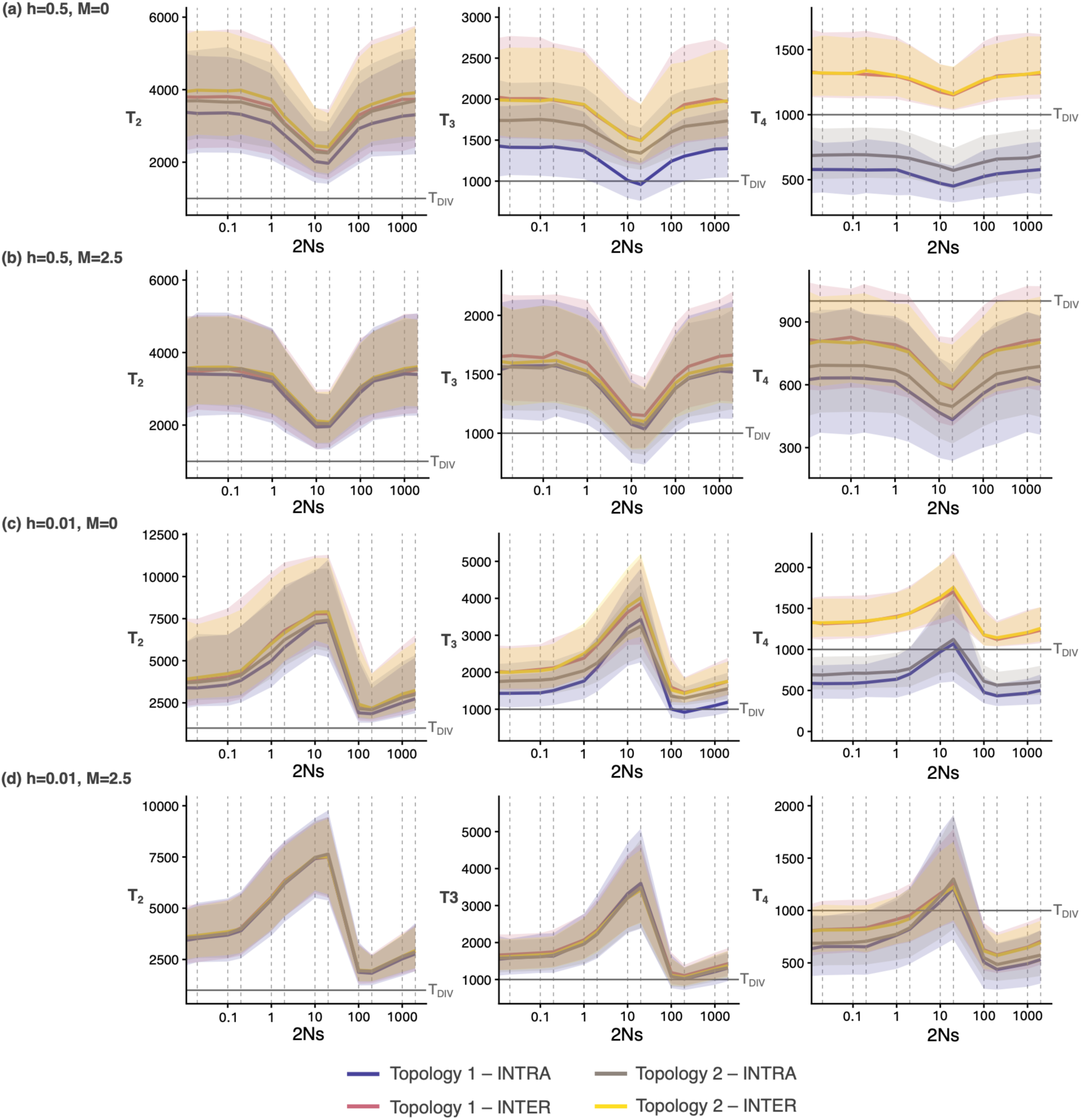
AOD inflates the coalescence times, while BGS compresses the coalescence times for codominant (a and b) or recessive mutations (c and d), in the absence (a and c) or presence of migration (b and d). For each panel, the left panel represents the median time to the last *coalescent* event (T2, i.e., T_MRCA_), the central panel the median time to the second coalescent event (T3) and the right panel the median time to the first coalescence event (T4). Colors represent the different topologies: T1_INTRA_ (blue); T1_INTER_ (red); T2_INTRA_ (grey); T2_INTER_ (yellow), the areas around the curves the 75% confidence interval computed over 1024 simulations. The vertical grey lines represent the 2Ns values used in the simulations and the horizontal grey line represent the divergence time T_DIV_. The x-axis is displayed in log scale.

## Discussion

Interpreting patterns of genetic variability along the genome is not trivial, as they result from a complex interaction between historical demography and selection. Here, we studied this interplay by conducting a simulation study, to quantify the effects of deleterious mutations on linked neutral variation in a structured scenario of an Isolation with Migration (**Figure 1**). Our study highlighted two remarkable outcomes. First, as expected, we found stronger effects of deleterious mutations on linked neutral variation for the lowest recombination rate investigated (i.e., *r* = 2.5e-8 = *µ* / 10 per site and per generation), recovering the relationship between genetic diversity and recombination predicted by analytical models (Charlesworth et al. 1993; Charlesworth et al. 1997; Charlesworth 2012). As the recombination rate varies greatly along the chromosomes (Cutter and Payseur 2013), we can expect that different regions of the genome will be differentially affected by linked selection, depending on local recombination rate. Second, as expected, we recover the same signatures on summary statistics when using *genotype*-based or *ARG*-based approaches, but the latter allowed a finer understanding of the effect of deleterious mutations on linked neutral regions through the direct study of gene genealogies.

### Transition from BGS to AOD in structured populations with migration

Our simulation results supported two regimes of variation in genetic patterns that depended on dominance and on the strength of selection of deleterious mutations. These are consistent with first a regime of associative overdominance (AOD), found for slightly deleterious (2*Ns*<20) recessive mutations (*h*=0.01), followed by a transition from AOD to BGS. These patterns are consistent qualitatively with the ones described by Zhao and Charlesworth (2016), Becher et al. (2020) and Gilbert et al. (2020). However, our results also matched quantitively the results from Gilbert et al. (2020), while being higher than the one reported by Zhao and Charlesworth (2016) or Becher et al. (2020) (respectively 2*Ns*=1 or 2.5 for the regime AOD). Such discrepancies are likely tied to the difference between approaches with Zhao and Charlesworth (2016) focusing on single large panmictic population and few loci, and Becher et al. (2020) simulating partially recessive mutations (*h*=0.2, based on *Drosophila melanogaster*). In contrast, Gilbert et al. (2020) assumed complete recessivity of deleterious mutations, a single population, with parameters tailored to humans, and further supported by an analytical multi-locus approach. Finally, we also detected a BGS-driven regime for moderately strong codominant mutations (2<2*N*s<200), consistent with theoretical predictions that deleterious mutations should not be too weak nor too strong to detect the effect of BGS (Charlesworth 2012).

Most characterizations of the effect of deleterious mutations have been derived for panmictic populations (Charlesworth et al. 1993; Charlesworth 2012; Zhao and Charlesworth 2016; Cvijović et al. 2018; Becher et al. 2020; Gilbert et al. 2020). Here, we devised a framework allowing us to understand the dynamics of BGS and AOD in structured populations. For codominant mutations (*h*=0.5), and in the absence of gene flow (*M*=0), we detected (1) decreases in diversity *(π* and *d_xy_)*, *DAFi*, and in the number of shared polymorphism; and (2) increases in *F_ST_* and in the number of private polymorphism (**Figures 2 and 3**). These patterns hold for strongly deleterious recessive mutations and are consistent with expectations of BGS, whereby the continual removal of linked deleterious mutations also removes linked neutral variation (Charlesworth et al. 1997; Nordborg 1997; Zeng and Corcoran 2015; Aeschbacher et al. 2017; Gilbert et al. 2020). For recessive weakly deleterious mutations, we observed another regime characterized by higher diversity (high *π, d_xy_*), higher *DAFi*, higher proportion of shared polymorphism and lower *F_ST_* than neutrality, consistently with AOD-driven regimes observed in panmictic populations (Zhao and Charlesworth 2016; Becher et al. 2020; Gilbert et al. 2020). This, along with the BGS-driven regime detected for higher selective coefficients than the AOD-driven regime, corroborates the evidence of the AOD-BGS continuum (Zhao and Charlesworth 2016; Becher et al. 2020; Gilbert et al. 2020) (**Figures 2 and 3**).

Migration alters the purging of deleterious mutations and thus also affects the genomic signatures of AOD and BGS. Relative to isolation, the main difference is that deleterious mutations are shared between the two populations, having two consequences: (1) no signatures of AOD nor BGS are detected on genetic differentiation, but (2) clear signatures of AOD and BGS remaining detectable on *π, π_g_, d_xy_* and *d_xy-g_,* the distortion of *b* values being of similar magnitude. As for *π*, the isolation scenario can be viewed as a proxy for the expected results of a panmictic scenario, since lineages sampled within a single population evolve without migration. Our results also support that *π_g_* is slightly more affected in scenarios with gene flow than in isolation and a single panmictic population scenarios (see **Table S3**). Altogether, our results suggest that AOD, BGS, and the transition between the two regimes remain detectable with gene flow, and that the combined behavior of summary statistics could distinguish between isolated and gene flow models. Importantly, these two regimes detected for recessive mutations mirror the AOD-BGS transition previously described in panmictic populations (Zhao and Charlesworth 2016; Gilbert et al. 2020), demonstrating that this continuum also applies in the case of a structured population both with and without gene flow.

### Deleterious mutations consistently affect gene genealogies

The effect of deleterious mutations on the coalescent process has been mostly investigated in panmictic populations. In this context, BGS is known to strongly decrease the coalescence times for codominant mutations (Hudson and Kaplan 1988; Nordborg 1997), whereas the consequences of recessive mutations on the properties of gene genealogies remain less explored. This bias is even more pronounced in structured populations, where the genealogical consequences of linked deleterious mutations has received little attention, except for codominant mutations (Zeng and Corcoran 2015). To address this gap in knowledge, we used an *ARG*-based approach by sampling a single diploid individual per population. By analyzing the sequence of local trees, we quantified the distribution of coalescence times associated with each topology and their relative proportions. Across genealogy-based statistics (topology and coalescence times), we consistently detected a BGS-driven regime for codominant deleterious mutations (*h*=0.5) and an AOD–BGS transition for recessive ones (*h*=0.01). Our results showed that the effects of BGS and AOD leave distinct signatures on the distribution of topology, and opposite effects on coalescent times, which can be simply described as a systematically shrinking under the BGS-driven regime and stretching under the AOD-driven regime. Importantly, these shifts in coalescent times are unaffected by population structure, while topology frequencies are affected by the interaction between gene flow and selection.

Following the “constraints” on coalescent events, topologies can be interpreted as proxy of population structure, with a decreasing level of structure following the ordering *T1_INTER_ > T2_INTER_ > T_2INTRA >_ T_1INTRA_*. These constraints can similarly be interpreted in terms of tree congruence, with the most congruent topology being *T_1INTRA_* and the least one, *T_1INTER_*. In isolation, BGS leads to a strong excess of the most structured topology and a strong deficit of the least structured topologies (*INTER-*deme topologies, **Figure 4a**). Going backwards in time, this pattern arises due to the increasing probability of removing a deleterious mutation with time, shifting coalescence times towards more recent values (**Figure 5**). This is consistent with theoretical work predicting that BGS leads to an increase in coalescence rate that can be interpreted as an effective reduction in *Ne* (Hudson and Kaplan 1988; Charlesworth et al. 1993). In structured populations, this will result in higher likelihood of a within-deme coalescent event than a between-deme coalescent event, which is only possible after *T_DIV_*. In contrast to BGS, AOD produces an excess of the two least structured topologies (*T1_INTER_* and *T2_INTER_*), and a deficit of *T1_INTRA_*. Under this regime, lineages sampled within the same deme carry complementary haplotypes. This stretches coalescence times (**Figure 5**); therefore, most topologies would come from situations where *T_4_* is older than *T_DIV_*. Consequently, the distribution of topologies should converge to the single population distribution of topologies for which *T2* topologies are twice as represented as *T1* topologies, which roughly matches the distribution of topologies at the AOD extremum (**Table S2**). Interestingly, we observe that removing the constraint of isolation erases all signatures of BGS on the distribution of topologies, but weak signatures of AOD remain detectable, suggesting a stronger robustness of AOD signatures to migration than BGS ones (but see limitations).

### From gene trees to summary statistics

The behavior of summary statistics (*π, d_xy_, F_ST_, SFS* and *DAF_i_*) systematically follows the variations detected in coalescence times and in topology proportions. This is expected given the duality between genealogy-based statistics and mutation-based summary statistics (Ralph et al. 2020; Wong et al. 2024).

In the BGS-driven regime, coalescence times are shrunk due to the removal of linked mutations. In isolation, between deme coalescence times (*INTER* topologies) are constrained by the divergence time (*T_DIV_*). This constraint leads to a stronger decrease in *π* than *d_xy_*, resulting in an increase in *F_ST_* relative to neutrality (Charlesworth et al. 1997; Cruickshank and Hahn 2014). With migration, this constraint is relaxed and within and between deme coalescence times are reduced by similar factors, leading to comparable reduction of *d_xy_* and *π* and to *F_ST_* values close to neutral values. The *SFS* and *DAFi* are respectively less and not impacted by gene flow. In this BGS-driven regime, shorter coalescence times and increase of *INTRA* topologies make it more likely to observe an excess of low frequency mutations and a deficit of shared ones between the two isolated demes in the *SFS* due to the absence of coalescence between lineages sampled in different demes. With gene flow, this effect remains detected, although less clearly, which suggests that coalescent times are sufficiently recent that deleterious mutations had a low probability of migrating between demes before the coalescence occurred. *DAF_i_* decreases similarly with and without gene flow, which is expected as the direct property of this statistic is to be sensitive to the effect of deleterious mutations irrespective of demography (Pouyet et al. 2018).

Under the AOD-driven regime, coalescence times are stretched because of the maintenance of haplotypes with different sets of slightly deleterious mutations. Irrespective of gene flow, this inflation directly leads to larger values of *π*, *d_xy_* and *DAF_i_* compared to neutrality, but *F_ST_* only decreases under isolation. These patterns can be explained within structured coalescent framework, noting that coalescent events are only possible between haplotypes with the same number and set of deleterious mutations. Going backwards in time, as lineages of neutral alleles move into haplotypes with less deleterious mutations, coalescent rates can differ at recent and older times depending on the number of deleterious mutations that were accumulated in the different haplotypes. With AOD and limited recombination, different haplotypes are likely to accumulate different sets of mutations, which increases coalescent times. Indeed, AOD affects more the recent portion of genealogies than the older one, due to the most complementary haplotypes being maintained among extant individuals. Therefore, the first coalescence events happen further into the past, resulting in an increase of *π*. Consequently, this unequal effect of AOD on coalescent time limits our ability to represent AOD as a simple a rescaling of the coalescent time. Such complementary haplotypes progressively break down going backwards in time, reducing the impact of AOD on older coalescent events, with the increase in coalescence times becoming progressively less pronounced. Consequently, as under isolation *d_xy_* remains constrained by *T_DIV_*, AOD has a weaker impact on *d_xy_* than on *π,* resulting in a reduction in *F_ST_* in comparison to neutrality. Under migration, *d_xy_* is no longer constrained by *T_DIV_* because gene flow continuously reshuffles individuals across demes. Thus, recent coalescence times within and between populations are similarly affected, leading to comparable increases in *π* and *d_xy_*, explaining why *F_ST_* values remain close to neutral expectations. Although the effect of AOD decreases with genealogical depth, coalescence among major complementary haplotypes remains delayed relative to neutrality. Consequently, deeper ancestral branches contribute more to the genealogy relative to the neutral expectations. Mutations arising on these branches are inherited by a larger number of descendants and therefore reach higher derived allele frequencies, inflating *DAFi* and shifting the *SFS* toward more shared variants. This shift is particularly noticeable under isolation where gene flow does not reshuffle alleles between populations and thus maintains the parallel sorting of haplotypes.

### Implications for inference of population genetics parameters

While theoretical, our study provides several key implications for inference of BGS and AOD in natural structured populations where empirical signatures consistent with these processes have been reported (Jackson et al. 2015; Torres et al. 2018; Becher et al. 2020; Gilbert et al. 2020; Leitwein et al. 2021; Waller 2021). Previous studies have showed that BGS can significantly bias estimates of population sizes (Ewing and Jensen 2016; Marsh and Johri 2024) and that AOD can produce false signatures of migrations under isolation (Smith and Hahn 2024). Here, we show that BGS and AOD systematically alter coalescent times, by shrinking and stretching genealogies, respectively, both with and without gene flow. In addition, we show that BGS and AOD alter the distribution of topologies in isolation but not with gene flow. This implies that linked deleterious mutations alone can generate substantial variations in coalescent times and in the shape of the genealogy (i.e., the distribution of topologies) across neutral regions in the genome (**Figure 6**), which has been shown to impact ARG-based inference (although in non-equilibrium panmictic populations (Marsh and Johri 2024)).

**Figure 6.**
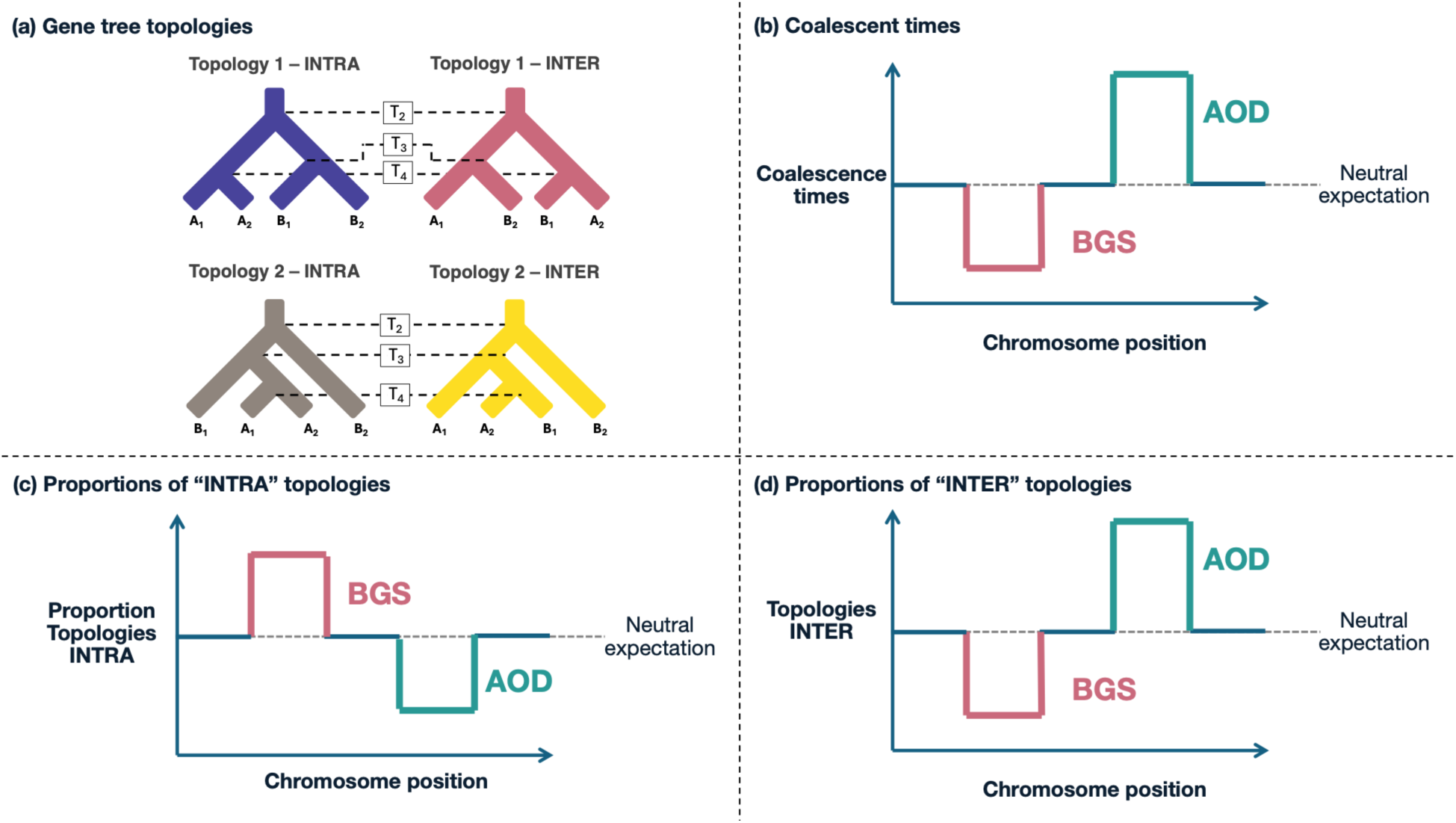
Schematic diagram representing the potential effect of AOD and BGS along the genome on different summary statistics. **(a)** The four topologies: T1_INTRA_ (blue) T1_INTER_ (red) T2_INTRA_ (grey), and T2_INTER_ (yellow). **(b)** The effect of BGS and AOD on coalescence times along the genome. **(c)** and **(d)** represent the effect of AOD and BGS on the proportion of topologies “INTRA” and “INTER”, respectively. The positions of the BGS and AOD regions along the chromosome are illustrative and do not imply that these effects depend on chromosomal position.

Second, because commonly used statistics such as *π*, *d_xy_*, *F_ST_* or the *SFS* are direct functions of coalescence times, they consequently yield signatures of BGS and AOD. These signals arise without changes in demographic parameters (*e.g.,* effective size, divergence time or migration rate), highlighting the risk of misinterpreting genomic heterogeneity.

Third, given (1) the distinct signatures on gene genealogies; (2) the direct relationship between summary statistics and gene genealogies; and (3) the ability to detect these signatures with one diploid sampled per population; our findings emphasize the potential of *ARG*-based inference to detect linked purifying selection in structured populations, although our study did not formally test the ability of using inferred *ARG*s to recover patterns of AOD and BGS. Although recent methods have been developed to detect selection across the genome in structured populations (Laetsch et al. 2023), none, to our knowledge, leverage explicit tree-based estimates and incorporate recessive mutations, highlighting the potential of such *ARG*-based approaches. We note that estimates based on *n*=2 diploids may exhibit great variance, but this could be minimized by aggregating multiple pairwise comparisons.

Finally, given the different effects of deleterious mutations in different recombination scenarios, our study suggests the ability to detect the effects of BGS and AOD by comparing regions of different recombination rates across the genome.

### Limitations and future directions

Despite the broad set of parameters explored, our study has several limitations. First, although the average recombination rate is usually of same order of magnitude of the mutation rate (Kong et al. 2010; Chan et al. 2012; Lipson et al. 2015) the effect of recombination rate could be further investigated, by considering intermediate values between our three recombination rates.

Secondly, our framework relies on fixed values of selection coefficients, whereas distributions of fitness effects are complex (Eyre-Walker and Keightley 2007) and often modeled using gamma or exponential distributions (Becher et al. 2020). To check the congruence of our results with this, we performed additional simulations drawing 2*Ns* values from an exponential distribution (mean *2Ns*=20) which recovered comparable signatures (**Table S4, Figure S6**). In addition, empirical studies in model organisms (*Drosophila*, *Arabidopsis*, *Saccharomyces*) suggest that dominance and strength of selection are usually correlated for deleterious mutations (Phadnis and Fry 2005; Huber et al. 2018; Becher et al. 2020). Our results with a fixed s=0.01 and h=0.01 were qualitatively similar to simulations where dominance and selection coefficients were drawn from a joint distribution following the formula of Huber et al. (2018), with an average s of 0.01. Indeed, the summary statistics are consistent with an AOD-driven regime, although with slightly weaker effects than in the corresponding single-value case (*s*=0.01, *h*=0.01, **Table S4**) and roughly similar to the case of drawing 2*Ns* values from an exponential distribution (mean 2*Ns*=20, i.e., *s*=0.01), with a fixed *h*=0.01. Although this should be further investigated in the future, these results indicate that the patterns detected under a simplified scenario capture major trends of more complex and realistic scenarios.

Third, the genealogical information generated by different dominance regimes suggests that ARG-based statistics may provide information on inferring dominance from genealogies in structured populations. In single populations it is difficult to estimate separately s and h, since predictions depend on their product, making s and h non-identifiable (Sendrowski et al. 2026). Yet, it has been shown that by comparing closely related outcrossing and selfing species it is possible to disentangle s from h (e.g., *Arabidopsis*, Huber et al. (2018)). Our results suggest that in structured populations it might be possible to disentangle s and h, as they lead to different gene trees. Yet, whether the ARG information is sufficient to jointly infer demography, selection and dominance remains to be tested.

Fourth, we modeled a simplified a genomic architecture corresponding to a single chromosome (**Figure 1**). While this choice allows for clear interpretation, it ignores the heterogeneity in recombination rates and of type of genomic elements observed along real genomes. In this light, investigating more complex architectures with heterogeneity in genomic elements and recombination rates will be an important next step.

Fifth, we mostly focused on the presence *versus* absence of migration. Exploring a wider range of migration values would allow to more quantitively investigate the effect of migration. To investigate in more detail the effect of migration rate, we investigated the case of *M*=0.5 (**Table S6**). We found that signatures of BGS were less pronounced than for *M*=0 but more visible than for *M*=2.5, especially in the distribution of topologies, suggesting the ability of the gene tree statistics to distinguish different demographic scenarios. Finally, the IM model investigated is a simplified proxy of population structure. Most species might display a more complex demographic history and population structure, such as stepping-stone scenarios, which should be investigated in the future. In addition, varying demographic parameters, such as effective sizes, divergence times and migration (including asymmetric) will be key to fully characterize the effects of BGS and AOD on summary statistics. Based on our results, we speculate that looking at the relative values of statistics (*b*), by comparing genomic regions under selection with neutral regions should remain affected similarly (i.e., *b*<1 or *b*>1) regardless of the demographic history, similar to statistics that are robust to demographic history such as *DAF_i_* (Pouyet et al. 2018).

Finally, we did not examine linkage disequilibrium statistics. Because BGS shortens coalescence times whereas AOD extends them, the two regimes may differ in how recombination decorrelates genealogies, resulting in distinct patterns of genealogical correlations and LD decay (**Figures S7, S8 and S9**). Testing this prediction is a natural extension of the current work.

## Conclusion

Our results show that the impact of deleterious mutations on linked neutral diversity in structured populations is strongly modulated by dominance, recombination, and gene flow. When recombination is low (0.1\**µ*), deleterious mutations affect linked neutral regions, thus leaving strong signatures on gene genealogies. We demonstrate that codominant deleterious mutations, produced a BGS-driven regime, while recessive deleterious ones produced two regimes: an AOD-driven regime directly followed by a BGS-driven regime (Zhao and Charlesworth 2016; Becher et al. 2020; Gilbert et al. 2020). BGS and AOD leave different and distinguishable patterns: BGS compresses coalescent times and increases the proportion of lineage sorting topologies, while AOD inflates coalescent times and shifts topology distribution in the opposite direction. We showed that these effects have a different behavior depending on gene flow. Finally, the fact that these signatures remain detectable from minimal sampling in an ARG-based framework highlights the potential of tree-based approaches to jointly infer selection, dominance and demography in structured populations.

## Data availability statement

All scripts to run simulations and estimate statistics are available in GitHub repositories: https://github.com/PierreLesturgie/Pipeline-Slim-BGS-AOD and https://github.com/PierreLesturgie/GTS and will be published on Zenodo upon acceptance)

## Acknowledgments

We thank Bárbara Parreira for her help in developing scripts. We also thank Gonçalo Caneira and João Frazão for their help with pilot study simulations.

## Author contributions

Conceptualization: P.L., V.C.S. Formal Analysis: P.L, A.B. Funding Acquisition: P.L, V.C.S. Investigation: P.L., A.B. Methodology: P.L. Validation: P.L., A.B., V.C.S. Visualization: P.L. Supervision: V.C.S. Writing—original draft: P.L. Writing—review and editing: P.L., A.B., V.C.S. All authors gave final approval for publication.

