## Supplementary figures for "The Joint Impact of Deleterious Mutations, Dominance, and Gene Flow on Linked Neutral Variation in Structured Populations"

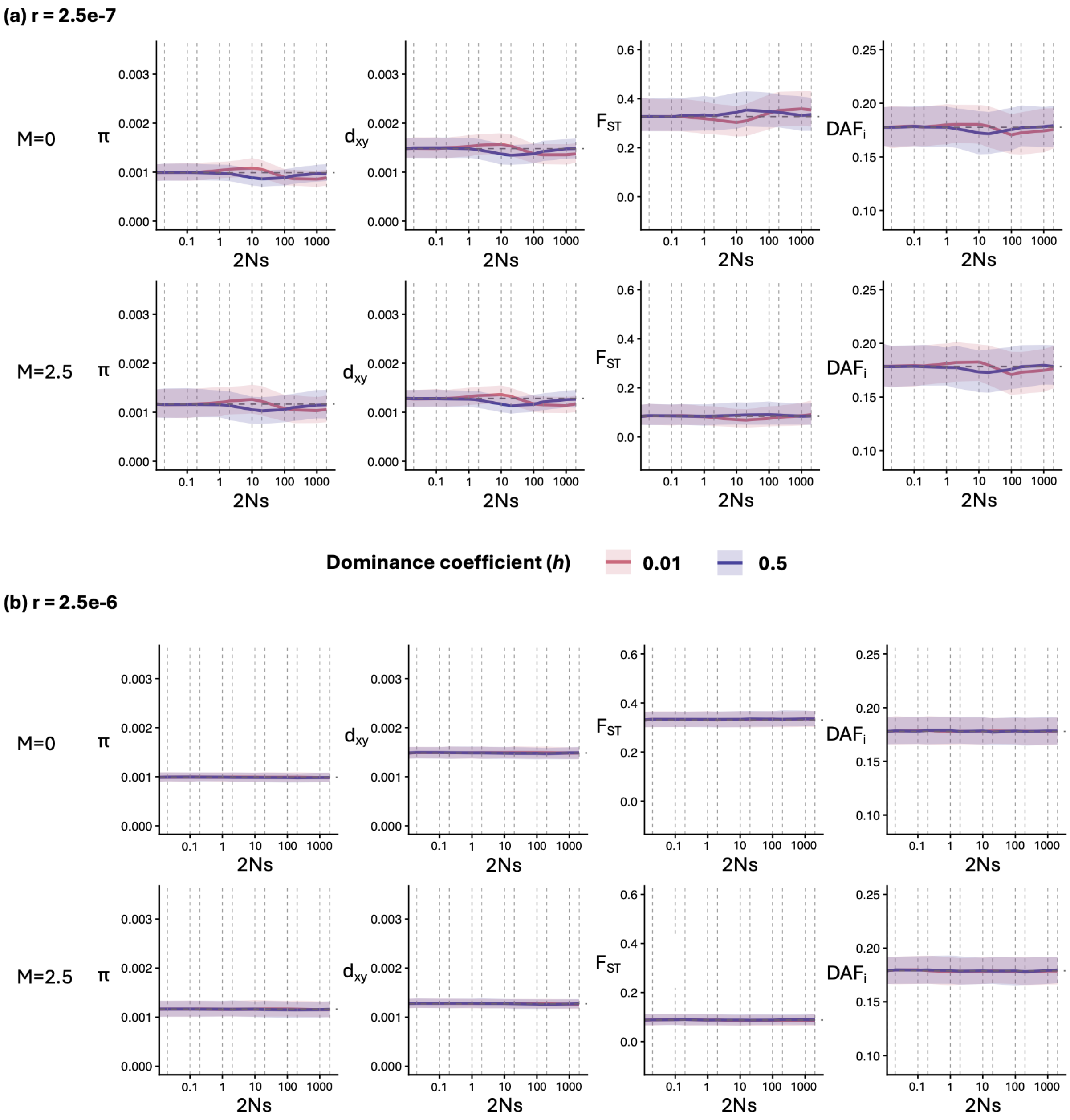


***Figure S1. Effect of the deleterious fitness effect on summary statistics computed using the genotype-based approach under recombination rates of r=2.5e-7 (a) and 2.5e-6 (b) per site per generation.*** *In each panel, the first row represents summary statistics computed under the no migration (M=0) scenario and the second the summary statistics computed under the scenario with migration (M=2.5). The summary statistics displayed are the mean pairwise difference (π), the absolute divergence ($d_{xy}$), the genetic differentiation ($F_{ST}$), and the Average Derived Allele Frequency per individual (DAFi). For each statistic, the trajectory of the median value as a function of the selective coefficient (2Ns) is represented and the areas around the curve represent*

*the 75% confidence interval computed over 1024 simulations. For each plot, the two curves represent co-dominant (h=0.5) mutations (blue) or recessive (h=0.01) mutations (red). The vertical grey lines represent the 2Ns values used in the simulations and the dashed horizontal line represent the median neutral value. The x-axis is displayed in log scale.*

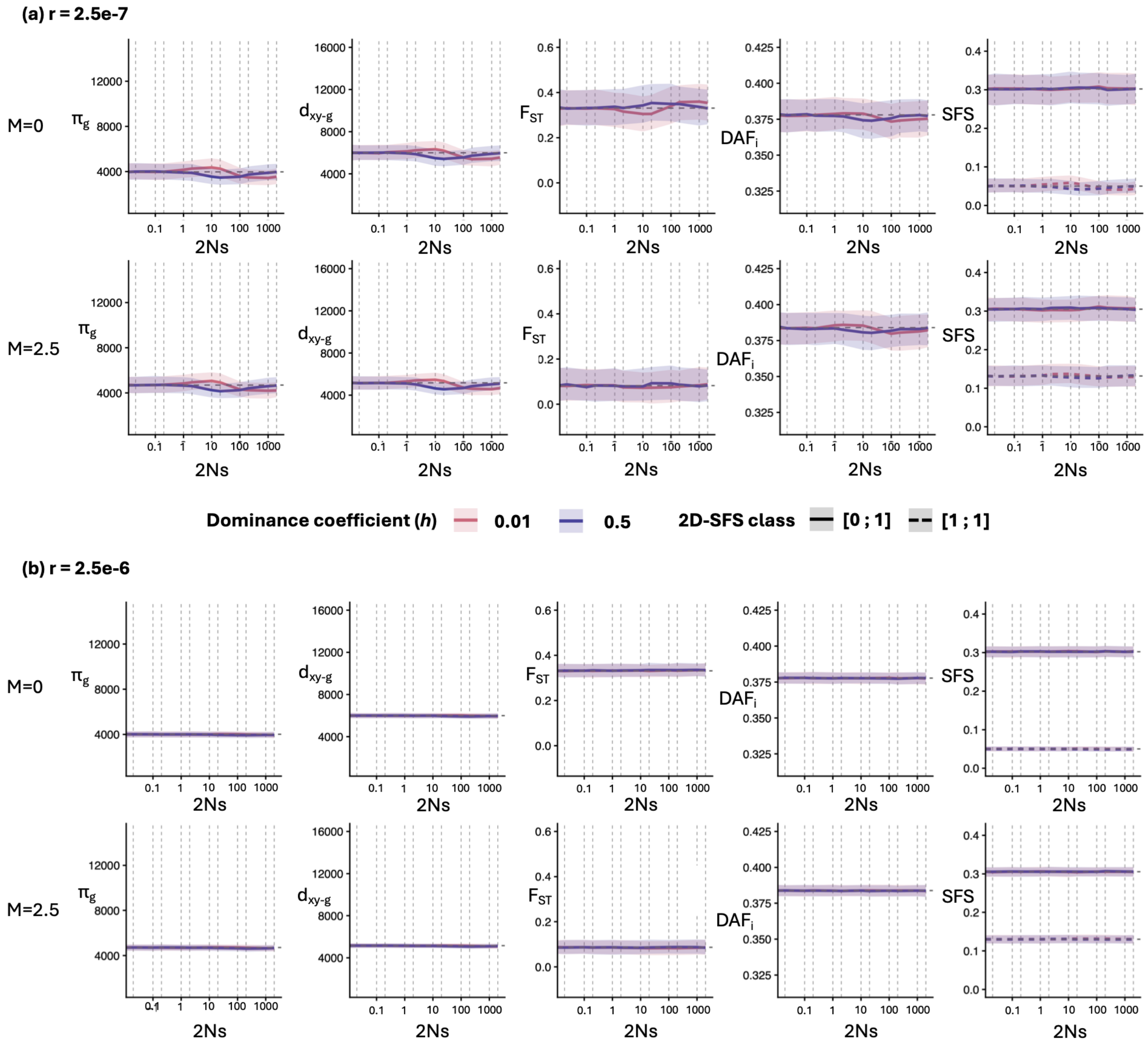


***Figure S2. Effect of the deleterious fitness effect on summary statistics computed using the ARG-based approach under recombination rates of r=2.5e-8 (a) and 2.5e-6 (b) per site per generation****. For each panel, the first row represents summary statistics computed under the no migration (M=0) scenario and the second row the summary statistics computed under the scenario with migration (M=2.5). The summary statistics displayed are the mean pairwise difference ($\pi_g$), the absolute divergence ($d_{xy-g}$), the genetic differentiation ($F_{ST}$), the Average Derived Allele Frequency per individual (DAFi) and the private (full line) or shared (dashed line) singletons classes of the two-dimensional Site Frequency Spectrum (2D-SFS). The $\pi_g$ and $d_{xy-g}$ are computed in branch lengths in generations and respectively represent $2*E[T_{within}]$ and $2*E[T_{between}]$, where $T_{within}$ and $T_{between}$ are the coalescent times for a pair of lineages sampled in the same population, or one from each population, respectively. For each statistic, the trajectory of the median value as a function of the selective coefficient (2Ns) is represented and the areas around the curve represent the 75% confidence interval computed over 1024 simulations. For each plot, the two curves represent co-dominant (h=0.5) mutations (blue) or recessive (h=0.01)*

*mutations (red). The vertical grey lines represent the 2Ns values used in the simulations and the dashed horizontal line represent the median neutral value. The x-axis is displayed in log scale.*

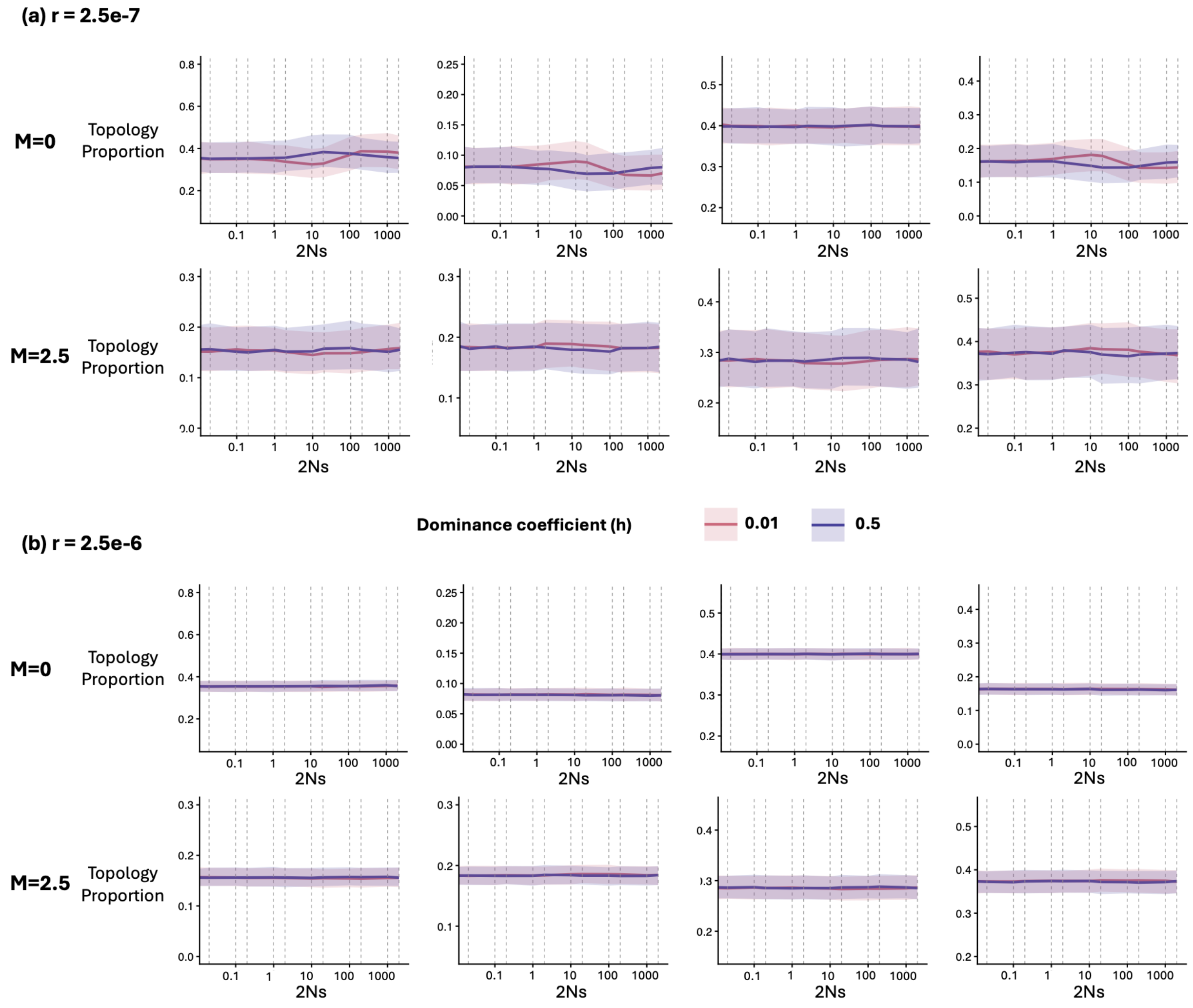


***Figure S3. Effect of the deleterious fitness effect on the distribution of topologies under recombination rates of r=2.5e-7 (a) and 2.5e-6 (b) per site per generation.*** *For each panel, the first row represents the distribution of topologies computed under the no migration (M=0) scenario and the second row the distribution of topologies computed under the scenario with migration (M=2.5). Considering four haploid genomes (A1 and A2 in population A; B1 and B2 in population B), Topology 1 is observed when the first and second coalescent events occur between external branches and Topology 2 happens when the first coalescent event occurs between two external branches and the second between the resulting internal branch and an external branch. Two subsets are devised: Topology 1 INTRA ($T1_{INTRA,}$ first column) and Topology 2 INTRA ($T2_{INTRA,}$ third column) occur when the first coalescent occurs between genomes from the same population. Topology 1 INTER ($T1_{INTER,}$ second column), and Topology 2 INTER ($T2_{INTER,}$ fourth column) occur when the first coalescent occurs between genomes from different populations. For each proportion of topology, the trajectory of the median value as function of the selective coefficient (2Ns) is represented and the areas around the curve represent the 75% confidence interval computed over 1024 simulations. For each plot, the two curves represent codominant (h=0.5) mutations (blue) or recessive (h=0.01) mutations (red). The*

*vertical grey lines represent the 2Ns values used in the simulations and the dashed horizontal line represents the median neutral value. The x-axis is displayed in log scale.*

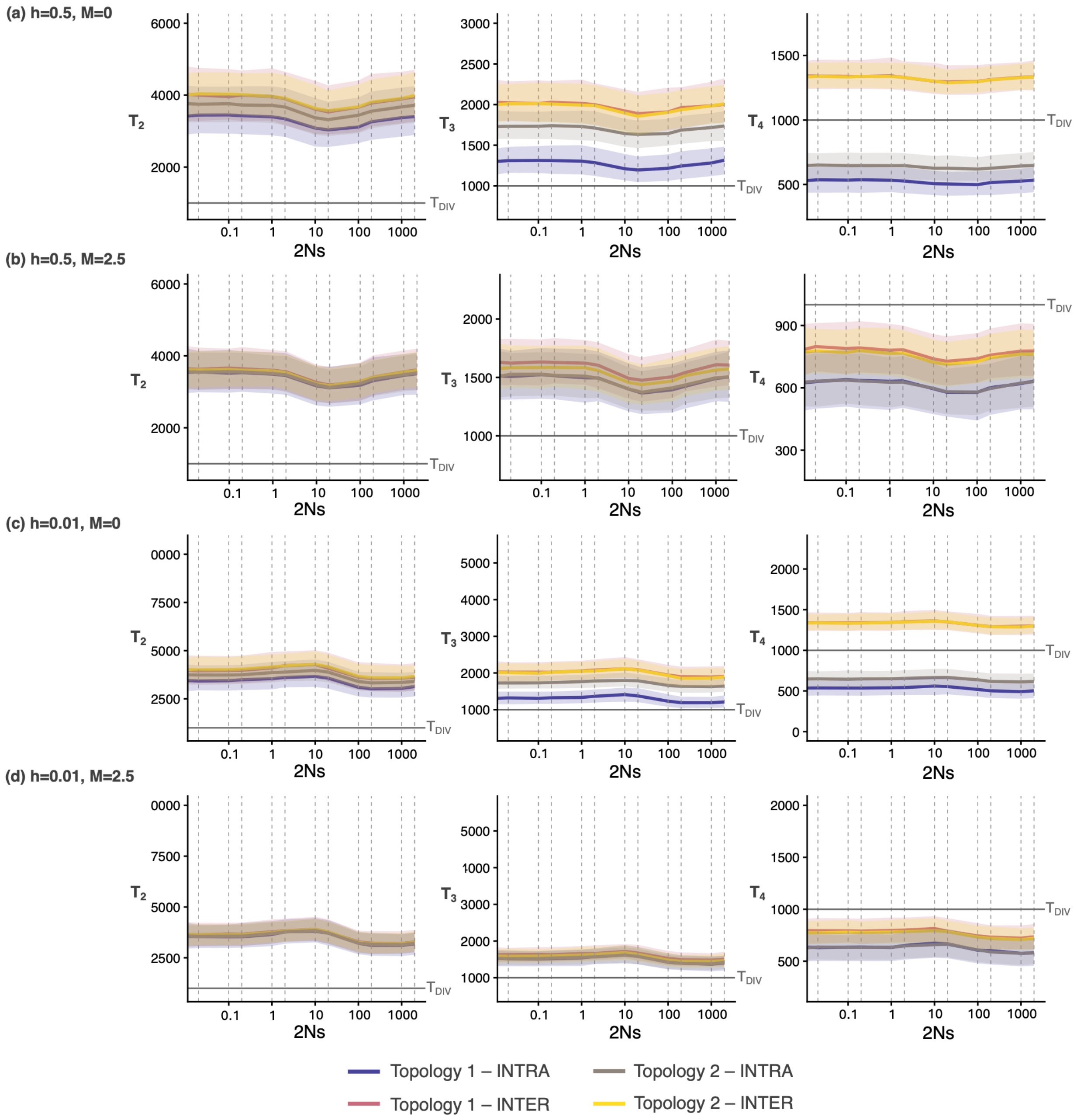


***Figure S4. Effect of the deleterious fitness effect on coalescence times under r=2.5e-7 per site per generation for codominant (a and b) or recessive mutations (c and d), in the absence (a and c) or presence of migration (b and d).*** *For each panel, the left panel represents the median time to the last coalescent event ($T_2$, i.e., $T_{MRCA}$), the central panel the median time to the second coalescent event ($T_3$) and the right panel the median time to the first coalescence event ($T_4$). Colors represent the different topologies: $T1_{INTRA}$ (blue); $T1_{INTER}$ (red); $T2_{INTRA}$ (grey); $T2_{INTER}$ (yellow), the areas around the curves the 75% confidence interval computed over 1024 simulations. The vertical grey lines represent the 2Ns values used in the simulations and the horizontal grey line represent the divergence time $T_{DIV}$. The x-axis is displayed in log scale.*

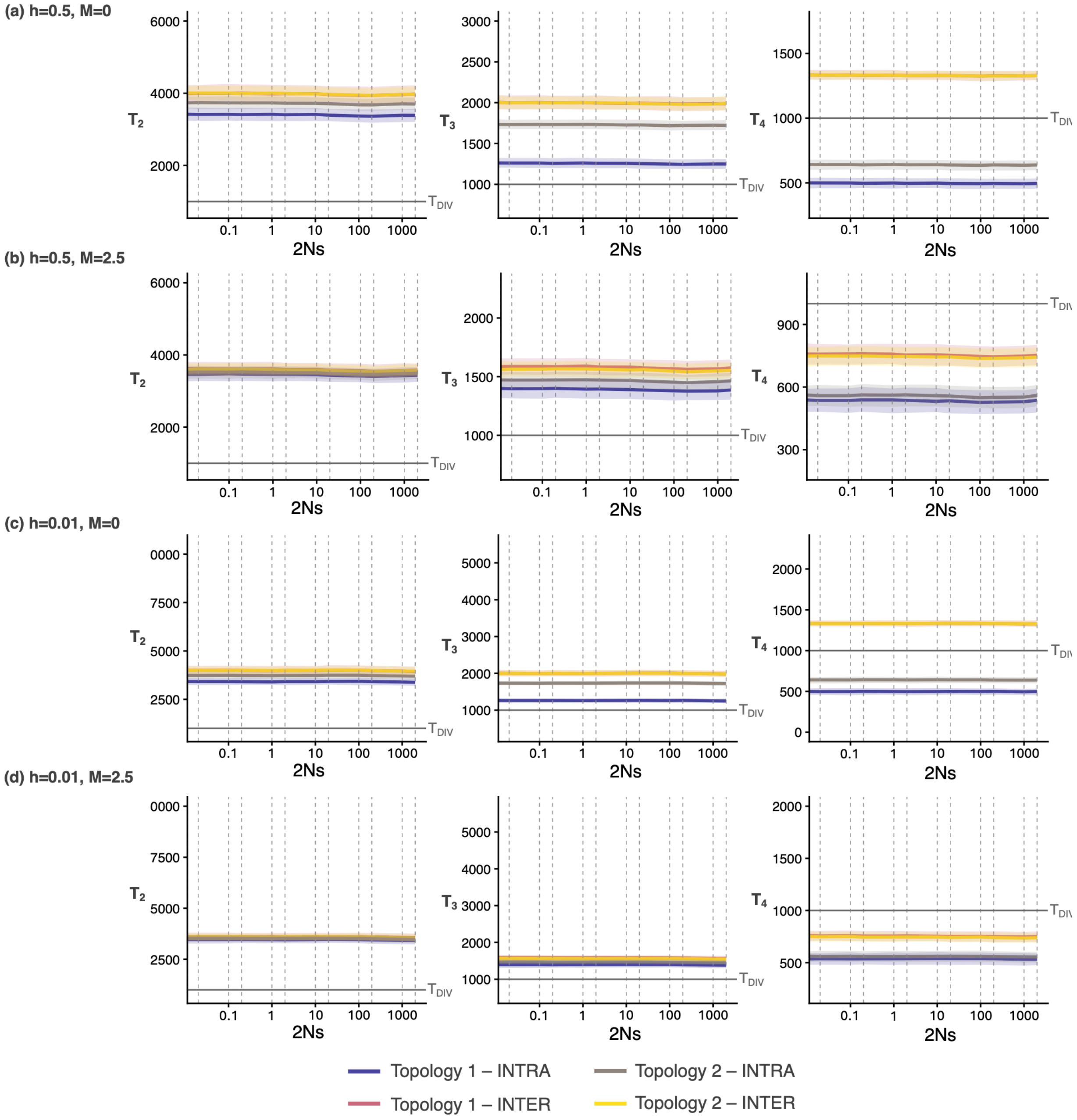


***Figure S5. Effect of the deleterious fitness effect on coalescence times under r=2.5e-6 per site per generation for codominant (a and b) or recessive mutations (c and d), in the absence (a and c) or presence of migration (b and d).*** *For each panel, the left panel represents the median time to the last coalescent event ($T_2$, i.e., $T_{MRCA}$), the central panel the median time to the second coalescent event ($T_3$) and the right panel the median time to the first coalescence event ($T_4$). Colors represent the different topologies: $T1_{INTRA}$ (blue); $T1_{INTER}$ (red); $T2_{INTRA}$ (grey); $T2_{INTER}$ (yellow), the areas around the curves the 75% confidence interval computed over 1024 simulations. The vertical grey lines represent the 2Ns values used in the simulations and the horizontal grey line represent the divergence time $T_{DIV}$. The x-axis is displayed in log scale.*

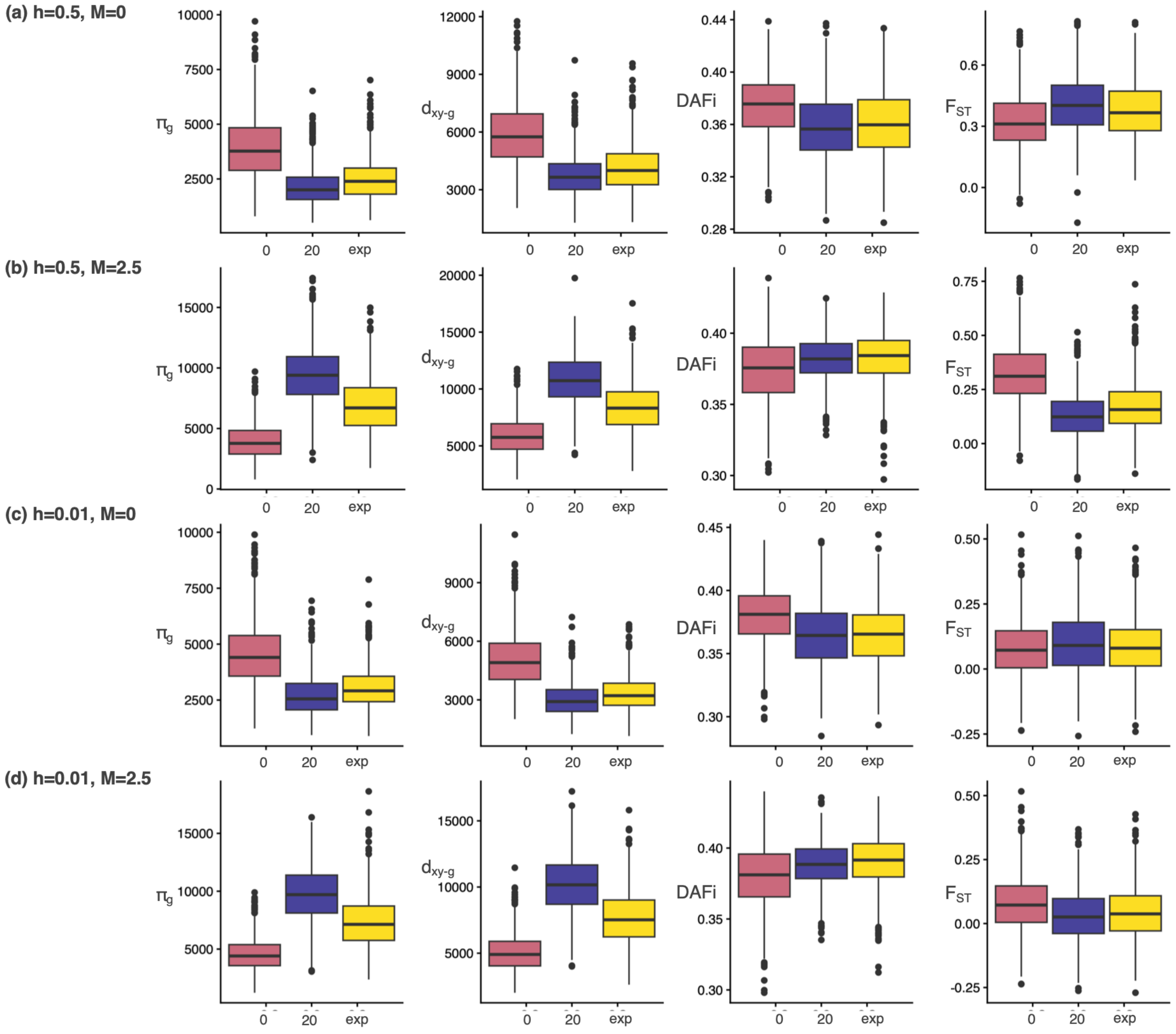


***Figure S6. Distribution of summary statistics under neutrality, under a constant 2Ns=20 and under an exponential distribution of mean 2Ns=20.*** ***(a)*** *Co-dominant mutations (h=0.5) under the isolation scenario (M=0).* ***(b)*** *Co-dominant mutations (h=0.5) under the gene flow scenario (M=2.5).* ***(c)*** *Recessive mutations (h=0.01) under the isolation scenario (M=0).* ***(d)*** *Recessive mutations (h=0.01) under the gene flow scenario (M=2.5). For each panel, the different plots represent different summary statistics:* $\pi_g$*; (first column),* $d_{xy\text{-}g}$*, (second column); DAFi, (third column) and* $F_{ST}$*, (fourth column) under neutrality (red boxplot); under a constant 2Ns=20 (blue) and under an exponential distribution of mean 2Ns=20 (yellow). The* $\pi_g$ *and* $d_{xy\text{-}g}$ *are computed in branch lengths in generations and respectively represent* $2*E[T_{within}]$ *and* $2*E[T_{between}]$*, where* $T_{within}$ *and* $T_{between}$ *are the coalescent times for a pair of lineages sampled in the same population, or one from each population, respectively.*

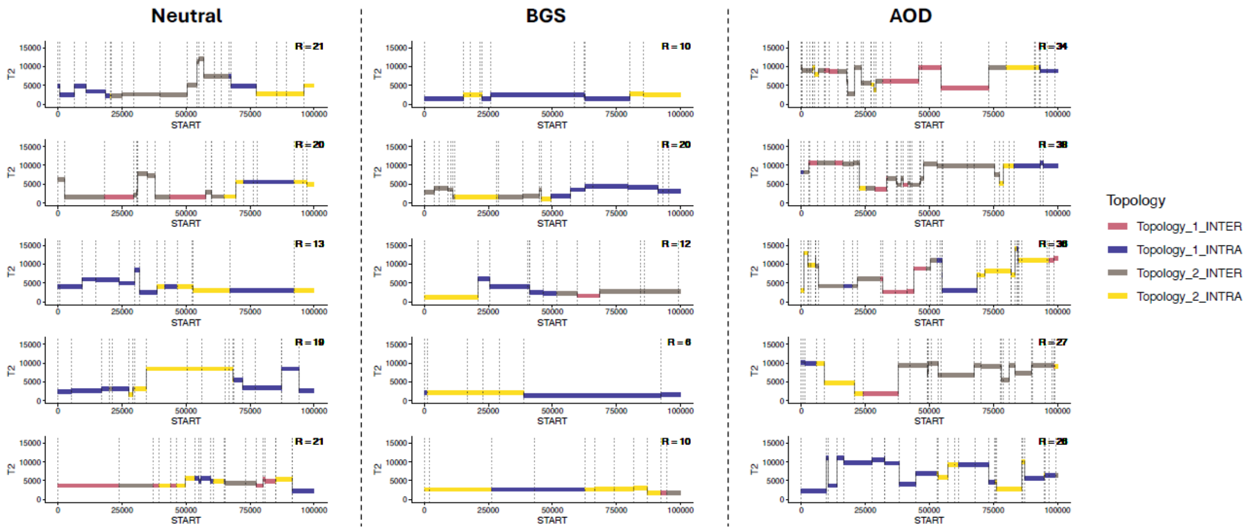


***Figure S7. Distribution of the TMRCA - time to the most recent common ancestor (T2) over the 100 Kb chromosome for the case with no migration (M=0) for five simulations***. *Each line corresponds to one simulation. The three columns correspond to a parameter combination corresponding to neutral (2Ns=0, h=0), BGS (2Ns=20, h=0.5) and AOD (2Ns=20, h=0.01). The vertical dotted lines correspond to recombination events that change the topology and/or coalescent times. Line colored according to the topology.*

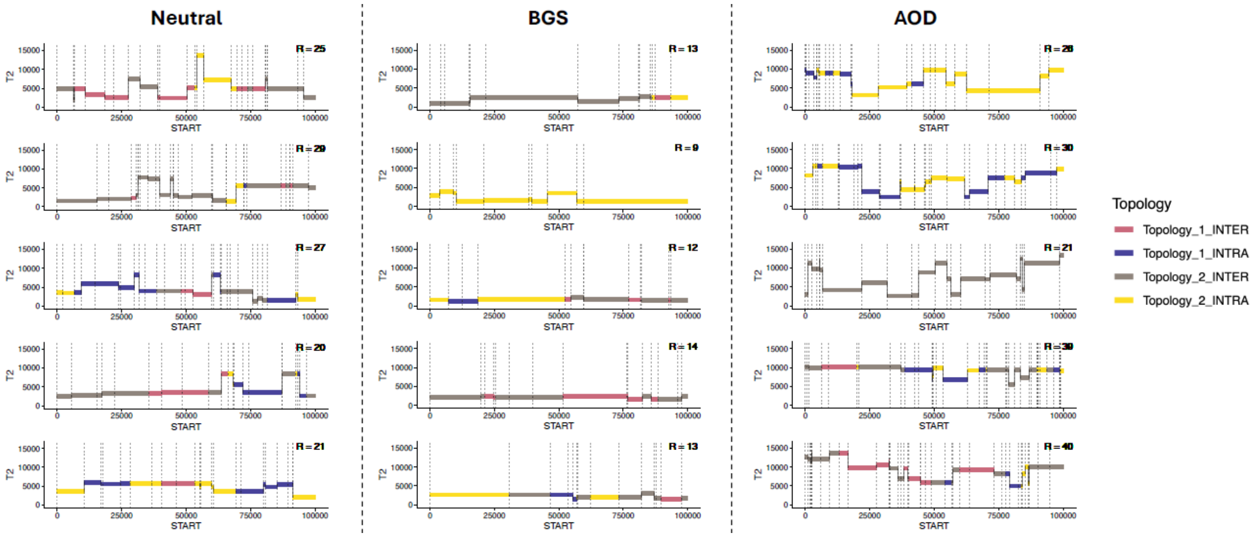


***Figure S8. Distribution of the TMRCA - time to the most recent common ancestor (T2) over the 100 Kb chromosome for the case with migration (M=2.5) for five simulations.*** *Each line corresponds to one simulation. The three columns correspond to a parameter combination corresponding to neutral (2Ns=0, h=0), BGS (2Ns=20, h=0.5) and AOD (2Ns=20, h=0.01). The*

*vertical dotted lines correspond to recombination events that change the topology and/or coalescent times. Line colored according to the topology.*

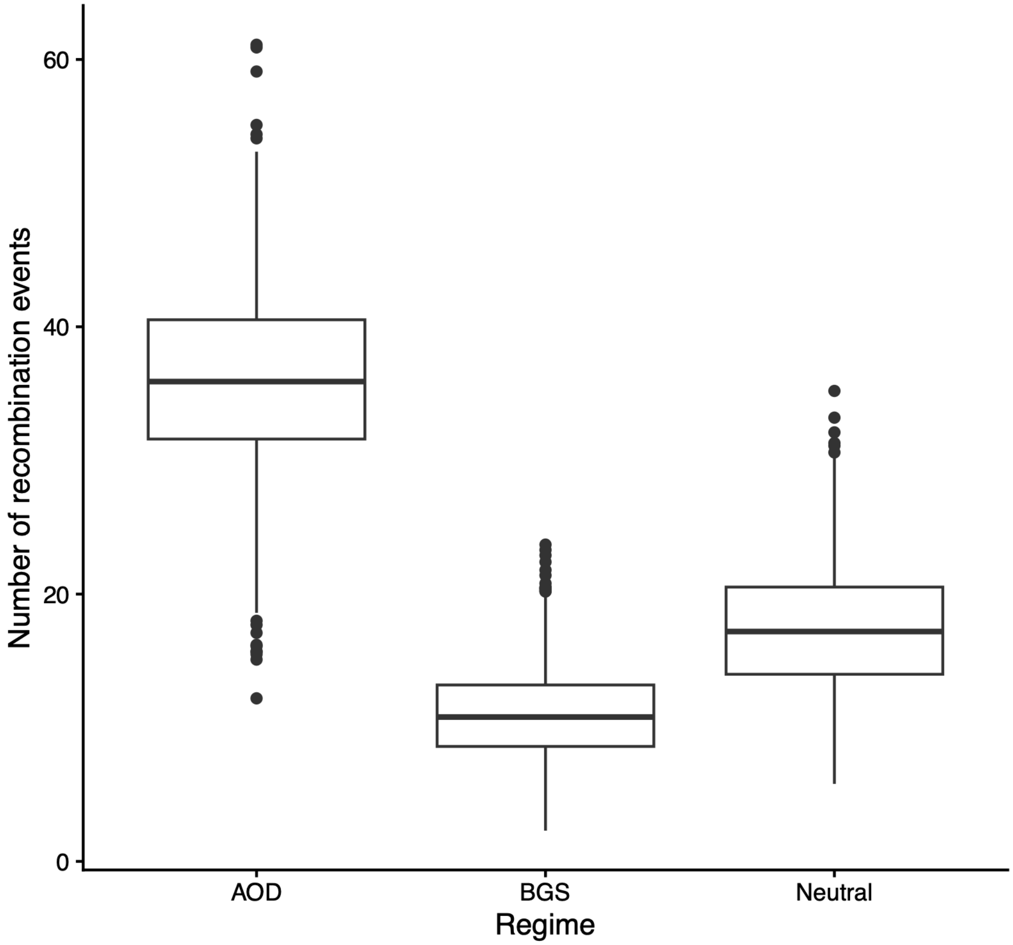


***Figure S9. Distribution of the recombination events under the AOD regime, BGS regime and Neutral regime.*** *Boxplots represent the distribution over the 1024 simulations.*
