## Supplementary material for "The Joint Impact of Deleterious Mutations, Dominance, and Gene Flow on Linked Neutral Variation in Structured Populations"

### Expected proportions of gene tree topologies in a model with no migration assuming neutrality

To validate if simulations under neutrality reflect theoretical expectations, we derived the proportions of the four gene tree topologies analytically. This is possible under a scenario without gene flow since we can enumerate all possible gene trees and compute their corresponding probabilities. Following the model used for the SLIM simulations, we consider a two population isolation model with an ancestral population that splits into two populations *TDIV* generations ago, all with the same effective population size (N). Moreover, we assume a sample of two lineages from population 1 (pop1) and two lineages from population 2 (pop2). To simplify, we distinguish coalescent events that happen in two epochs: (E1) epoch E1, when there is only one ancestral population, i.e., for coalescent times older than *TDIV*; (E2) epoch E2, when there are two descending populations, i.e., for coalescent times younger than *TDIV*.

Given that during epoch E2 the two populations evolve in isolation without migration, we can treat the coalescent within each population as independent Poisson processes. We assume a continuous time approximation with time is measured in units of *2N* generations. Hence, the probability of no coalescent events during *T* generations for a sample of two lineages from a population of size *N* is approximately exponential: $(1-(1/(2N)))^T \sim \text{Exp}[-(T/(2N))]$, where Exp represents the exponential function. In the continuous time approximation, the time of divergence is $t_d = TDIV/2N$.

Denoting the lineages sampled in population 1 as A, and lineages sampled in population 2 as B, there are four possible topologies (Fig. 4):

- T1 Intra: ((AA),(BB))
- T1 Inter: ((AB),(AB))
- T2 Intra: ((B,AA),B) or ((A,BB),A)
- T2 Inter: ((A,AB),B) or ((B,AB),A)

Looking backwards in time, for a sample of four lineages two from population 1 and two from population 2, i.e., AABB, we need to consider the following possibilities:

- Two coalescent events happen in Epoch E2, resulting in topology T1 Intra ((AA),(BB))
- One coalescent event happens in Epoch E2, all other coalescent events happen in Epoch E1, resulting in topologies T1 Intra [((AA),(BB))] or T2 Intra [((B,AA),B) or ((A,BB),A)].
- No coalescent events in Epoch E2, and all coalescent events in Epoch E1, resulting in all possible topologies, depending on the order of coalescent events in ancestral population.

There are two key probabilities we must consider:

- No coalescent in Epoch E2 for a given population *j*, for *j*=(1,2)

$$\Pr(\text{No coalescent in epoch D for pop } j) = \Pr(\text{No coal } j) = \text{Exp}[-t_d]$$

- Coalescent in Epoch E2 for a given population $j$, for $j$=(1,2)

  Pr(Coalescent in epoch D for pop $j$) = Pr(coal $j$) = 1-Exp[-$t_d$]

Once the lineages are in the ancestral population we need to consider either there are 4 or 3 lineages. Noting that (n chooses 2)= (n*(n-1))/2 denotes the number of pairs of lineages when there are n lineages left, if there are four lineages, there are (4 chooses 2)=6 possible pairs of lineages, and when there are three lineages there are (3 chooses 2)=3 possible pairs. For each case, we need to count of many of those are consistent with a given topology.

The following Table represents all events, corresponding probabilities and resulting topology. To obtain the probability we need to multiply for each row the probability of Epoch E2 with the probability of Epoch E1. Rows resulting in the same topology are summed.

| **Events Epoch E2** | **Probability Epoch E2** | **Events Epoch E1** | | **Probability Epoch E1** | **Resulting topology** |
|---|---|---|---|---|---|
| | | **4 lineages** | **3 lineages** | | |
| Two coalescent events in E2 | $(1-Exp[-t_d])^2$ | - | - | 1 | T1 intra ((AA),(BB)) |
| One coalescent event in E2 in pop1 | $(1-Exp[-t_d])$* $Exp[-t_d]$ | - | coalescent of two pop2 lineages BB | 1/3 | T1 intra ((AA),(BB)) |
| | | | coalescent of lineages of pop2 with pop1 B,AA | 2/3 | T2 intra ((B,AA),B) |
| One coalescent event in E2 in pop2 | $Exp[-t_d]$* $(1-Exp[-t_d])$ | - | coalescent of two pop1 lineages AA | 1/3 | T1 intra ((AA),(BB)) |
| | | | coalescent of lineages of pop1 with pop2 A,BB | 2/3 | T2 intra ((A,BB),A) |
| No coalescent event in E2 | $Exp[-t_d]^2$ | coalescent of two pop1 lineages AA | coalescent of pop2 lineages BB | (1/6)*(1/3) | T1 intra ((AA),(BB)) |
| | | | coalescent of lineages of pop2 B with AA | (1/6)*(2/3) | T2 intra ((B,AA),B) |
| | | coalescent of two pop2 lineages BB | coalescent of pop1 lineages AA | (1/6)*(1/3) | T1 intra ((AA),(BB)) |
| | | | coalescent of lineages of pop1 A with BB | (1/6)*(2/3) | T2 intra ((A,BB),A) |
| | | coalescent of lineages of the two pops AB | coalescent of pop1 and pop2 lineages AB | (4/6)*(1/3) | T1 inter ((AB),(AB)) |
| | | | coalescent of lineages of one pop with AB, i.e., A,AB or B,AB | (4/6)*(2/3) | T2 inter ((A,AB)B) or ((B,AB)A |

Replacing for *TDIV*=1000 generations, we obtain the following probabilities

| **Events Epoch E2** | **Probability Epoch E2** | **Probability Epoch E1** | **Probability with $t_d$=0.5** | **Resulting topology** |
|---|---|---|---|---|
| Two coalescent events in E2 | $(1\text{-Exp}[-t_d])^2$ | 1 | 0.155 | T1 intra ((AA),(BB)) |
| One coalescent event in E2 in pop1 | $(1\text{-Exp}[-t_d])$* $\text{Exp}[-t_d]$ | 1/3 | 0.080 | T1 intra ((AA),(BB)) |
| | | 2/3 | 0.159 | T2 intra ((B,AA),B) |
| One coalescent event in E2 in pop2 | $\text{Exp}[-t_d]$* $(1\text{-Exp}[-t_d])$ | 1/3 | 0.080 | T1 intra ((AA),(BB)) |
| | | 2/3 | 0.159 | T2 intra ((A,BB),A) |
| No coalescent event in E2 | $\text{Exp}[-t_d]^2$ | (1/6)*(1/3) | 0.020 | T1 intra ((AA),(BB)) |
| | | (1/6)*(2/3) | 0.041 | T2 intra ((B,AA),B) |
| | | (1/6)*(1/3) | 0.020 | T1 intra ((AA),(BB)) |
| | | (1/6)*(2/3) | 0.041 | T2 intra ((A,BB),A) |
| | | (4/6)*(1/3) | 0.082 | T1 inter ((AB),(AB)) |
| | | (4/6)*(2/3) | 0.164 | T2 inter ((A,AB)B) or ((B,AB)A |
| TOTAL | | | 1 | |

For each topology, these result in the following probabilities for TDIV=1000 generations, rounded to 2 decimal points, matching the proportions obtained in simulations.

| **Topology** | **Probability for TDIV=1000 generations** | **Proportion obtained in SLIM simulations for M=0 and s=0** |
|---|---|---|
| T1 intra | 0.35 | 0.35 |
| T1 inter | 0.08 | 0.08 |
| T2 intra | 0.40 | 0.40 |
| T2 inter | 0.16 | 0.16 |
